# Identifying a novel role for the master regulator *Tal1* in the Endothelial to Hematopoietic Transition

**DOI:** 10.1101/2021.10.03.462906

**Authors:** Yasmin Natalia Serina Secanechia, Isabelle Bergiers, Matt Rogon, Christian Arnold, Nicolas Descostes, Stephanie Le, Natalia Lopez Anguita, Kerstin Ganter, Chrysi Kapsali, Lea Bouilleau, Aaron Gut, Auguste Uzuotaite, Ayshan Aliyeva, Judith Zaugg, Christophe Lancrin

## Abstract

Recent progress in the generation of bona-fide Hematopoietic Stem and Progenitor Cells (HSPCs) *in vitro* and *ex vivo* has been built on the knowledge of developmental hematopoiesis, underscoring the importance of understanding in detail this developmental process. Here, we sought to elucidate the function of the hematopoietic regulators *Tal1*, *Lmo2* and *Lyl1* in the Endothelial to Hematopoietic Transition (EHT), the process through which HSPCs are generated from endothelial precursors during embryogenesis. We used a mouse embryonic-stem cell (mESC)-based differentiation system to model hematopoietic development, and combined gain-of-function experiments in sorted vascular smooth muscle cells (VSM) with multi-omics to obtain mechanistic insights into the mode of action of *Tal1*, *Lmo2* and *Lyl1*. We found that these factors promote the silencing of the VSM transcriptional program and the activation of the hematopoietic one. Through this approach and the use of a Tet-on system to control the expression of Tal1 during hematopoietic specification from mESCs, we discovered that its expression in endothelial cells is crucial for the EHT to occur.

## INTRODUCTION

An important goal of regenerative medicine is the production of Hematopoietic Stem and Progenitor Cells (HSPCs) *in vitro* or *ex vivo* for clinical applications. Remarkable proofs-of-concept have been achieved in recent years exploiting current knowledge on the developmental processes and molecular regulators of developmental hematopoiesis (Lis *et al*., 2017; Sugimura *et al*., 2017). This highlights the importance of gaining a deep understanding of this biological process to design effective regenerative approaches.

During embryonic life, HSPCs are produced transiently within the major embryonic arteries from specialized endothelial cells known as Hemogenic Endothelium (HE) (Bruijn *et al*., 2000; Lancrin *et al*., 2009). This event is conserved across vertebrate species, and has been identified as a transdifferentiation termed “Endothelial to Hematopoietic Transition” (EHT) (Bertrand *et al*., 2010; Boisset *et al*., 2010; Kissa and Herbomel, 2010).

The aim of our work was to understand the contribution of *Tal1*, *Lmo2* and *Lyl1* to the generation of pre-hematopoietic stem and progenitor cells (Pre-HSPCs), which represent the intermediate stage of the EHT (Bergiers *et al*., 2018). *Tal1*, *Lmo2* and *Lyl1* form with *Runx1*, *Gata2*, *Cbfb*, *Erg* and *Fli1* a group of eight master regulators the combined activity of which regulates HSPC development (Nicola K. Wilson *et al*., 2010; Guibentif *et al*., 2017; Bergiers *et al*., 2018). Using a mouse embryonic-stem-cell (mESC)-based hematopoietic differentiation model, we have previously identified two opposing forces within these factors, which determine whether cells retain an endothelial identity (*Erg* and *Fli1*) or undergo EHT (*Runx1*, *Cbfb* and *Gata2*) (Bergiers *et al*., 2018). The simultaneous co-expression of all eight factors maintains cells in the Pre-HSPC stage, but the exact role of *Tal1*, *Lmo2* and *Lyl1* in this context remained unclear (Bergiers *et al*., 2018). *Tal1* is essential for the onset of hematopoiesis and is required for the generation of the HE, but its expression in endothelial cells is considered dispensable for the EHT to take place (Robb *et al*., 1995; Shivdasanl, Mayer and Orkin, 1995; Porcher *et al*., 1996; Endoh *et al*., 2002; D’Souza, Elefanty and Keller, 2005; Schlaeger *et al*., 2005; Lancrin *et al*., 2009). Indeed, the conditional ablation of *Tal1* in endothelial cells did not impair mouse hematopoiesis (Schlaeger *et al*., 2005) and its conditional re-expression in *Tal1^-/-^* mESCs-derived cultures could not rescue hematopoiesis after endothelial cells were formed, which led to conclude that *Tal1* exerts its essential function before endothelial cell formation to prime the cells towards a hematopoietic fate, but does not have a prominent role at later stages (Endoh *et al*., 2002). In the first study, however, *Tal1* could have been expressed for sufficient time for HE to be formed. Conversely, in the second study the lack of TAL1 should have impaired HE generation, likely explaining why its re-expression at later stages could not rescue hematopoietic cell formation. LMO2 is a scaffold protein and a TAL1-binding partner, and its role in embryonic hematopoiesis mostly mirrors that of TAL1 (Schmeichel and Beckerle, 1994; Valge-Archer *et al*., 1994; Wadman *et al*., 1997; Yamada *et al*., 2000; Stanulovic *et al*., 2017). However, while *Tal1* ablation from mESCs completely abrogates hematopoiesis *in vitro*, *Lmo2* ablation is permissive to the generation of Pre-HSPCs and leads to later defects, suggesting that it is required for Pre-HSPC maturation (D’Souza, Elefanty and Keller, 2005; Lancrin *et al*., 2009; Stanulovic *et al*., 2017). LYL1 is a bHLH protein highly homologous to TAL1 in the bHLH domain (Mellentin, Smith and Cleary, 1989; McWilliam *et al*., 2013; Consortium *et al*., 2020). Possibly owing to this, *Lyl1* was able to compensate for the loss of *Tal1* in adult HSCs (Souroullas *et al*., 2009). However, it could not rescue hematopoiesis *in vitro* from *Tal1^-/-^* mESCs (Capron *et al*., 2006; Chan *et al*., 2006). The relevance of this protein for hematopoietic development is elusive, as *Lyl1*^-/-^ mice displayed only mild hematopoietic defects, while the generation of *Lyl1^-/-^* mice with an 80% reduction in *Tal1* expression in the erythroid compartment revealed a functional redundancy between *Lyl1* and *Tal1* in primitive erythropoiesis (Chiu *et al*., 2018). *Tal1*, *Lmo2* and *Lyl1* are co-expressed at the single-cell level in Pre-HSPCs (Bergiers *et al*., 2018) and *Tal1* overexpression in mouse yolk-sac endothelial cells increased their hematopoietic output, suggesting a role for it in the EHT (Kim *et al*., 2013).

In the present work, combining gain-of-function experiments to multi-omics approaches *in vitro* we found that *Tal1*, *Lmo2* and *Lyl1* promote the downregulation of the VSM transcriptional program and the activation of the hematopoietic one. Remarkably, using a Tet-on system to control the expression of *Tal1* in our mESC-based hematopoietic differentiation system, we showed for the first time that *Tal1* expression in the endothelium is essential for the generation of Pre-HSPCs.

## RESULTS

### The simultaneous overexpression of *Runx1*, *Cbfb*, *Gata2*, *Erg* and *Fli1*, but not that of *Tal1*, *Lmo2* and *Lyl1* generates a hemangioblast culture enriched in cells resembling Pre-HSPCs

To investigate the role of *Tal1*, *Lmo2* and *Lyl1* in the EHT, we exploited a mESC-based differentiation system that models hematopoietic development *in vitro* (Keller *et al*., 1993). The differentiation system and the cell-surface markers used throughout this study to identify and isolate the different cell populations in culture are described in Fig.1A. In our previous work (Bergiers *et al*., 2018) we had shown that the doxycycline (dox)-induced simultaneous overexpression of *Tal1*, *Lmo2* and *Lyl1* with *Erg*, *Fli1*, *Runx1*, *Cbfb* and *Gata2* (8TFs) (Fig.1B) gave rise to mESC-derived hematopoietic cultures (i.e. “hemangioblast cultures”) highly enriched in Pre-HSPCs. To identify the contribution of *Tal1*, *Lmo2* and *Lyl1* to this phenotype, we generated two additional mESC lines using the inducible cassette exchange method previously described, which can overexpress *Tal1*, *Lmo2* and *Lyl1* alone (3TFs) and *Runx1*, *Cbfb*, *Gata2*, *Erg* and *Fli1* (5TFs) in a dox-inducible manner (Fig. 1B). Differentiation of the new cell lines confirmed that both were able to generate the *in vitro*-equivalents of VSM (VE-CAD^-^CD41^-^), endothelial (Endo, VE-CAD^+^CD41^-^), Pre-HSPCs (VE-CAD^+^CD41^+^) and hematopoietic progenitor cells (HP, VE-CAD^-^CD41^+^) after 3 days of hemangioblast culture, as assessed by morphological analysis and flow cytometry (Fig. 1C). The overexpression of the 3TFs and 5TFs was induced by treatment with dox at day 1 of hemangioblast culture, and the cultures were analyzed by flow cytometry two days after treatment. We found that just like the individual overexpression of the 3TFs (Bergiers *et al*., 2018), their simultaneous overexpression did not have an obvious impact on the composition of the culture (Fig. 1C). In stark contrast, the overexpression of the 5TFs generated a culture highly enriched in Pre-HSPCs (Fig. 1C), similar to the overexpression of the 8TFs (Bergiers *et al*., 2018).

**Figure 1:**
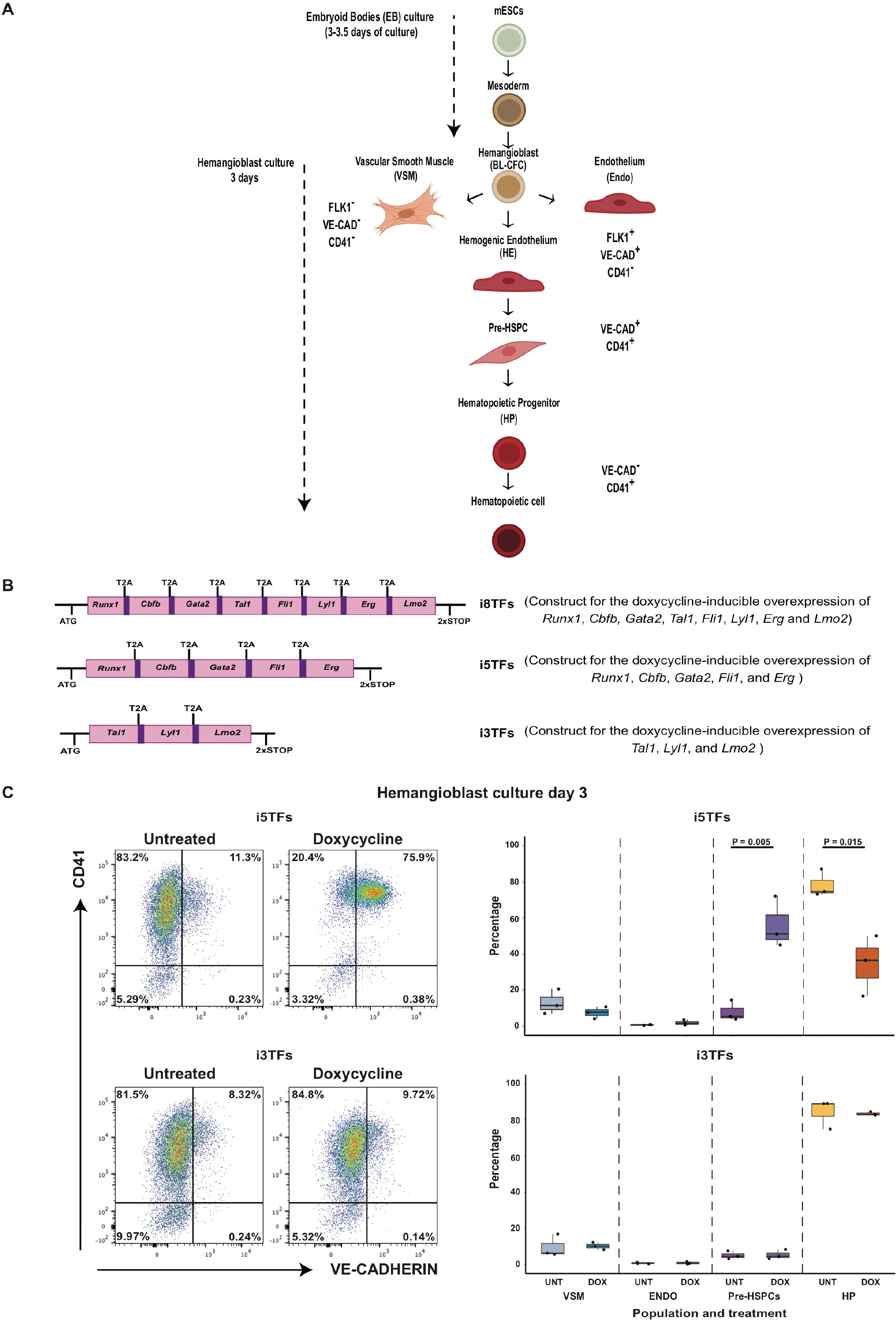
The simultaneous overexpression of *Runx1*, *Cbfb*, *Gata2*, *Erg* and *Fli1*, but not that of *Tal1*, *Lmo2* and *Lyl1* generates a hemangioblast culture enriched in cells resembling Pre-HSPCs. A. Scheme of the mESC differentiation system used in the study showing the cell-surface markers used to identify the different cell populations. B. Schemes describing the transgenes used in the study. T2A = viral T2A peptide sequences inserted between the coding sequences of the genes to exploit the “ribosomal skipping” mechanism for the production of all proteins from one single transcript. C. FACS analysis results showing the expression of VE-CAD and CD41 in day 3 hemangioblast cultures from i5TFs and i3TFs mESCs treated with doxycycline at day 1 of culture. Left: representative FACS plots. Right: Box plots summarizing the results of 3 experiments. Significance was determined by Analysis of Variance (ANOVA) test. Error bars correspond to standard deviations.

### The simultaneous overexpression of the 5TFs transdifferentiates sorted eVSM cells into VE-CAD^+^CD41^+^ Pre-HSPC-like cells less efficiently compared to the 8TFs

Through careful examination of the effect of 5TFs overexpression in hemangioblast cultures, we noticed the presence of a residual VSM population (Fig.1C) which was instead almost absent in cultures where the 8TFs were overexpressed (Bergiers *et al*., 2018). As *Tal1* has been reported to act as a repressor of cardiac fate in mesodermal and endothelial cells (Schoenebeck, Keegan and Yelon, 2007; Ismailoglu *et al*., 2008; Handel *et al*., 2012; Org *et al*., 2015; Chagraoui *et al*., 2018), this difference could reflect a role of the 3TFs also in repressing the VSM identity. We decided therefore to use an alternative assay to investigate the changes induced by differential transcription-factor overexpression. We chose a population enriched in VSM cells (eVSM) (Fig.2), because they provide a non-hematopoietic and non-endothelial background against which to test the activity of transcription factors. Moreover, they can be transdifferentiated into VE-CAD^+^CD41^+^ cells that resemble Pre-HSPCs by the simultaneous overexpression of the 8TFs (Fig. 1A & 2C) (Bergiers *et al*., 2018). As schematized in Fig.2A, we differentiated mESCs from the i8TFs, i5TFs and i3TFs cell lines and FACS sorted from each a population of FLK1^-^CD41^-^ eVSM cells (parent eVSM) at day 1 of hemangioblast culture (Fig.2A,C). Sorted FLK1^-^CD41^-^ cells were re-plated in a hemogenic endothelium (HE) medium that promotes hematopoiesis (Lancrin *et al*., 2009) in the absence or presence of dox to induce the overexpression of the transcription factors (Fig.2A). The cells were harvested after 1.5 days of HE culture and analyzed by flow cytometry (Fig. 2A, 2C & 2D). The overexpression of the 8TFs recapitulated previously published results, with the majority of the cells in culture co-expressing VE-CAD and CD41 (Fig. 2C & 2D). Differently from the hemangioblast culture, the overexpression of the 5TFs in sorted eVSM generated a culture enriched in VE-CAD^+^ cells, a large fraction of which did not co-express the hematopoietic cell-surface marker CD41 (Fig. 2C & 2D). Instead, similar to the hemangioblast culture, the overexpression of the 3TFs alone did not obviously alter the expression pattern of CD41 and VE-CAD in cultured cells compared to untreated controls, with the majority of the cells in culture displaying a VSM morphology (Fig. 2B, 2C and 2D). The overexpression of the 5TFs alone was therefore less efficient compared to that of all 8TFs in promoting the transdifferentiation of eVSM cells into Pre-HSPC-like cells. Thus, even though the 3TFs were not sufficient on their own to alter the phenotype of eVSM cells, they seemed to increase the ability of the 5TFs to promote the acquisition of a Pre-HSPCs phenotype in non-hematopoietic cells, which suggests that they might have important non-redundant roles in the generation of this cell type.

**Figure 2:**
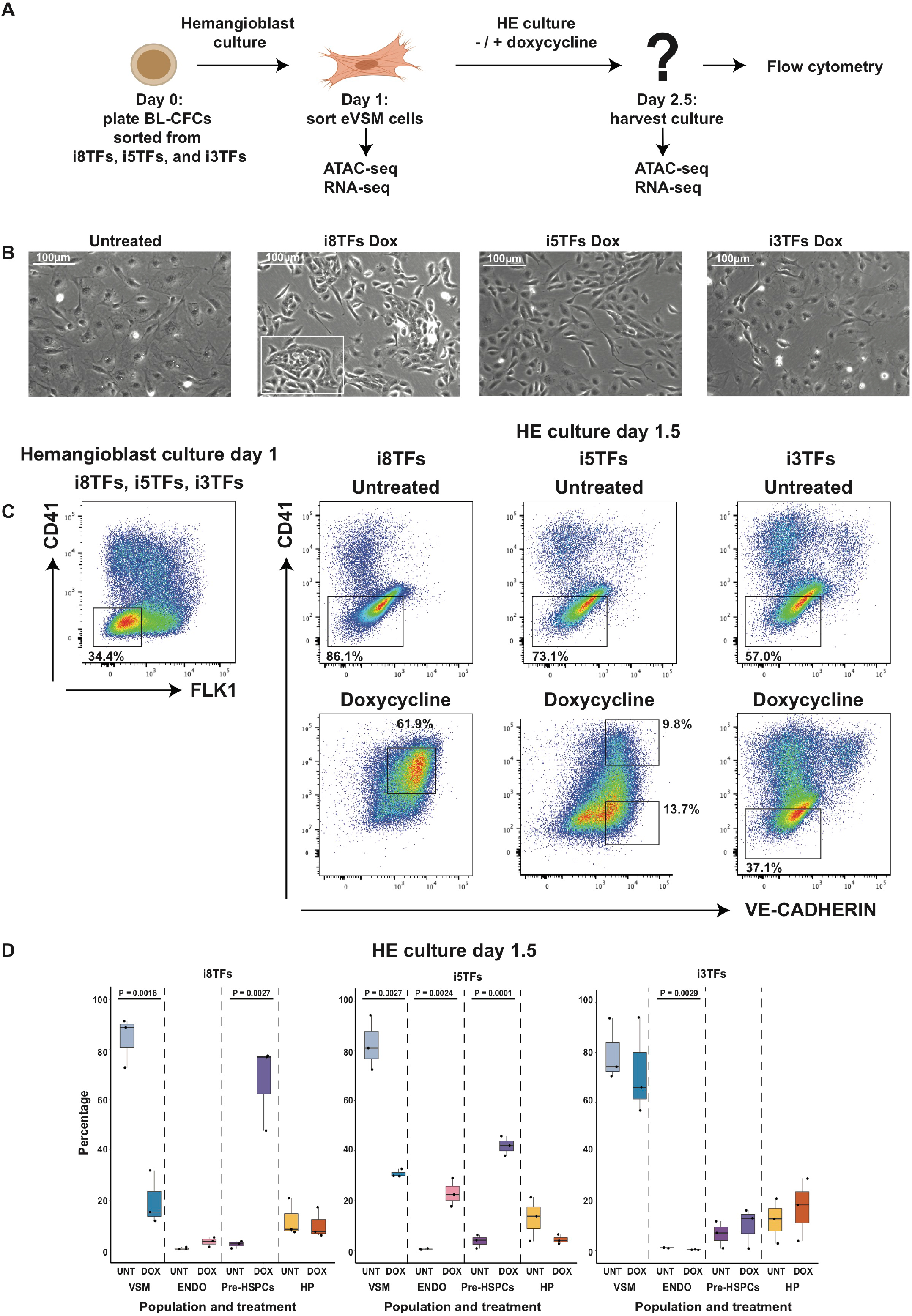
The simultaneous overexpression of the 5TFs transdifferentiates sorted eVSM cells into VE-CAD^+^CD41^+^ Pre-HSPC-like cells less efficiently compared to the 8TFs. A. Experimental layout of the work. B. Representative microscopy images of the indicated conditions at day 2.5 HE culture. VSM cells appear as large wide-spread cell; Endo cells appear as elongated cells; HPs appear as round floating cells. The white box highlights an example of the endothelial cell cluster from which hematopoietic cells normally arise. C. Flow cytometry analyses of the indicated conditions. The squares highlight the populations sorted for cell culture and molecular biology. D. Box plots showing the relative frequencies of VSM, Endo, Pre-HSPC and HP populations in HE culture from i8TFs, i5TFs and i3TFs lines (n=3). Significance was determined by Analysis of Variance (ANOVA) test. Error bars correspond to standard deviations.

### 8TFs overexpression in eVSM led to a higher number of chromatin regions being closed compared to 5TFs overexpression

To get a mechanistic insight on the changes induced by differential transcription factor overexpression in eVSM cells, we FACS sorted the most abundant cell types that were generated in the three dox-treated conditions as well as VE-CAD^-^CD41^-^ eVSM (late eVSM) cells from untreated controls, and assessed their chromatin accessibility and transcriptional profile by ATAC-seq and RNA-seq (Fig. 2A & 2C) (Picelli *et al*., 2014; Buenrostro *et al*., 2015). The sorting gates are shown in Fig. 2C. Principal component analysis (PCA) on the chromatin accessibility and transcriptional data (Fig.3A) confirmed that the parent eVSM sorted from the three cell lines were qualitatively similar, as they clustered together in both analyses. Cells in which the 8TFs and 5TFs were overexpressed greatly differed from untreated controls in their chromatin accessibility and transcriptional landscapes, while cells where the 3TFs were overexpressed appeared to be overall similar to untreated controls, in line with the morphological and cytofluorimetric observations.

**Figure 3:**
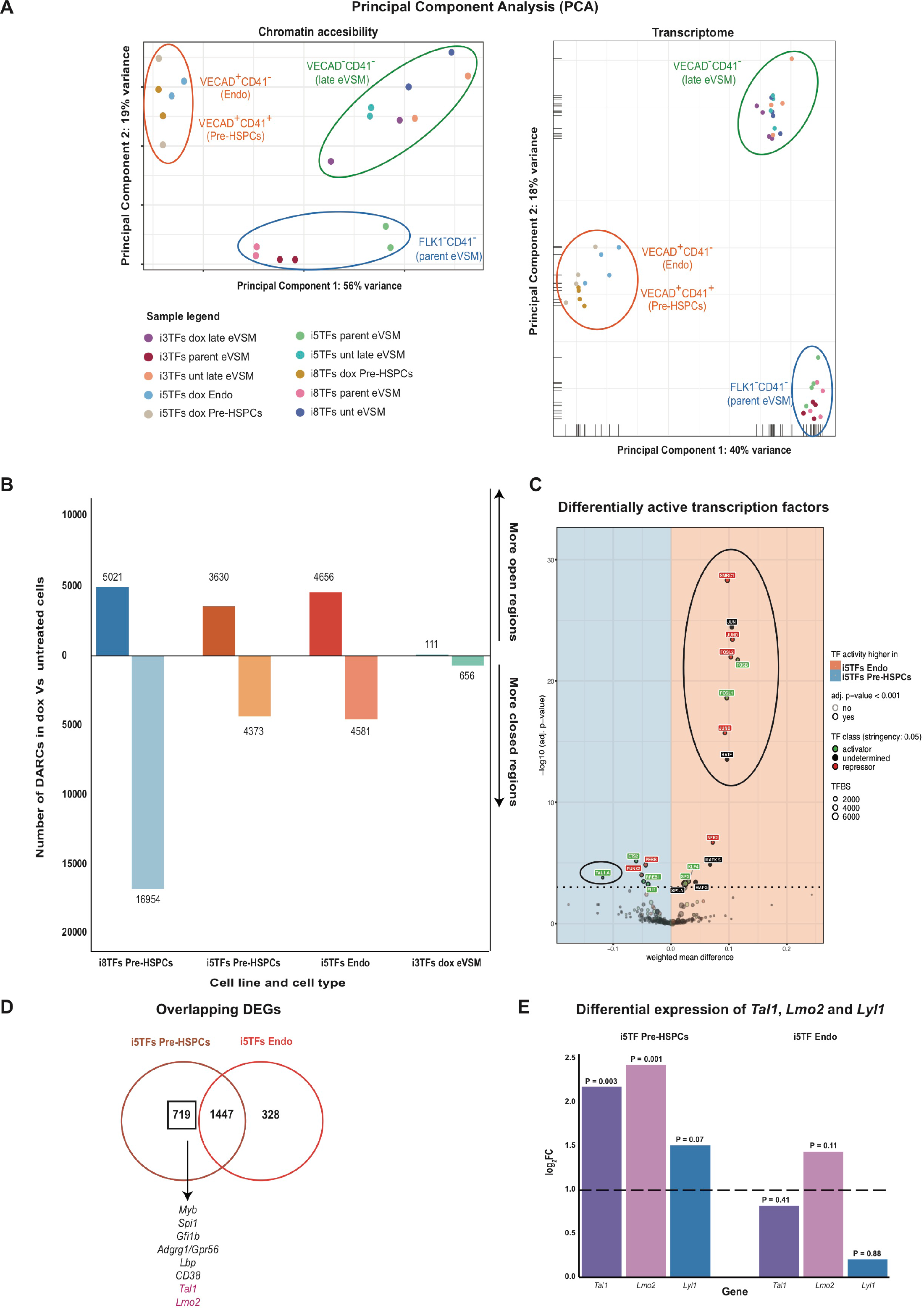
The endogenous *Tal1* and *Lmo2* genes were differentially expressed in i5TFs Pre-HSPCs compared to i5TFs Endo cells. A. PCA plots of ATAC-seq results (n=2) (left panel) and RNA-seq (n=4) results (right panel). Parent eVSM= FLK1^-^CD41^-^ cells sorted at day 1 of hemangioblast culture; late eVSM= VE-CAD^-^ CD41^-^ cells sorted at day 1.5 of HE culture; Endo= VE-CAD^+^CD41^-^ cells sorted at day 1.5 of HE culture; pre-HSPCs= VE-CAD^+^CD41^+^ cells sorted at day 1.5 of HE culture; dox= cultured in HE medium with doxycycline; unt= cultured in HE medium without doxycycline. B. Bar plots showing the number of chromatin regions identified by DiffBind analysis to be more accessible (more open) and less accessible (more closed) in the i8TFs pre-HSPCs, i5TFs pre-HSPCs, i5TFs Endo and i3TFs dox VSM cells compared to untreated controls. C. Results of diffTF analysis for the indicated cell lines. TFs identified as more active in i5TFs Pre-HSPCs are displayed in the blue quadrant, TFs identified as more active in i5TFs Endo are displayed in the red quadrant. TFs classified as activators are labeled in green, TFs classified as repressors are labeled in red, differentially active TFs that couldn’t be clearly classified as either are labeled in black. 5% of TFs were classified as activators or repressors (TF class stringency: 0.05) based on the Pearson correlation index. The x axis (weighted mean difference) shows the difference in TF activity between the two conditions. The y axis displays the significance of the TFs. The significance threshold is indicated with a dotted line (FDR adjusted p-value < 0.05). TFs are indicated as a dot. The size of each dot is proportional to the number of predicted genomic TFBS for each (TFBS). D. Venn diagrams comparing the DEGs in i5TFs Pre-HSPCs and i5TFs Endo. A subset of hematopoietic genes differentially expressed in Pre-HSPCs but not Endo cells is shown. E. Bar plots showing the differential expression (log2FC) of *Tal1*, *Lmo2* and *Lyl1* in i5TFs pre-HSPCs and i5TFs Endo cells compared to untreated eVSM cells, identified by RNA-seq.

We then used the DiffBind package (Stark and Brown, 2011) to identify the regions of chromatin that differed in their accessibility profile in dox-treated cells compared to untreated controls. We identified 767 differentially accessible regions of chromatin (DARCs) between untreated and dox-treated i3TFs eVSM cells (Fig. 3B), the majority of which were regions that were more closed in dox-treated cells. Interestingly, we found a substantial difference in the number of regions that became more closed upon 8TFs and 5TFs overexpression (Fig. 3B), as overexpression of the 8TFs resulted in the closure of approximately 17,000 regions of chromatin, whereas less than 5,000 regions of chromatin were more closed in cells where the 5TFs were overexpressed compared to untreated eVSM cells. A comparable number of chromatin regions was instead more open in cells where the 8TFs and the 5TFs were overexpressed (Fig. 3B). Thus, not only was the overexpression of the 5TFs less efficient in generating Pre-HSPCs compared to the overexpression of the 8TFs, but the cells generated were also different at the chromatin accessibility level. The overexpression of the 3TFs seemed to affect particularly chromatin closure, an effect that was much more pronounced when the 3TFs were overexpressed in combination with the 5TFs in the i8TFs cell line (Fig. 3B).

### DARCs seem to reflect the loss of VSM identity and the suppression of non-hematopoietic cell fates

To understand the biological meaning of the observed chromatin changes, we performed Gene Ontology (GO) analysis on the DARCs (Figure 3-figure supplement 1). As the majority of the DARCs mapped to intergenic regions, we used the Genomic Regions Enrichment Annotations Tool (GREAT) to perform the analysis on genes located within a defined distance from the DARCs. The majority of the biological processes terms enriched in regions of chromatin that were more open in 8TFs and 5TFs-overexpressing cells compared to untreated controls were related to hematopoiesis and the immune system, consistent with the nature of the tested factors as hematopoietic regulators (Figure 3-figure supplement 1B). Compared to 5TFs-overexpressing cells, we found in i8TFs Pre-HSPCs an enrichment in terms related to the SMAD signaling pathway, which has reported roles in regulating blood development, whereas terms related to the regulation of the p38 MAPK signaling pathway were enriched specifically in i5TFs Endo (Wilkinson *et al*., 2009; Souilhol *et al*., 2016). The two significant terms that could be identified for regions of chromatin more open in i3TFs dox eVSM belonged to the category “Mouse Phenotype Single Knock-out” and were related to erythropoiesis. Many of the GO terms enriched in regions of chromatin that were more closed compared to untreated controls appeared to reflect the loss of VSM identity (Figure 3-figure supplement 1C), as they referred to the organization of the extracellular matrix and cytoskeleton remodeling. We also found in i8TFs pre-HSPCs and i5TFs Pre-HSPCs terms related to TGFβ signaling, and in i8TFs pre-HSPCs the term “regulation of catenin import to the nucleus” related to the Wnt signaling pathway, both pathways which are involved in the regulation of the EHT (Ruiz-Herguido *et al*., 2012; Chanda *et al*., 2013; Souilhol *et al*., 2016; Vargel *et al*., 2016). The GO terms enriched in regions of chromatin that became more closed upon 3TFs overexpression were for the most part related to organ and tissue development, including gliogenesis and pericardium development, which might reflect a role of the 3TFs in suppressing non-hematopoietic developmental programs.

### The endogenous *Tal1* and *Lmo2* genes were differentially expressed in i5TFs Pre-HSPCs compared to i5TFs Endo cells

The cells generated by the induction of the 5TFs in eVSM cells differed in the expression of the hematopoietic cell-surface marker CD41 (Fig. 2C & 2D), suggesting that the hematopoietic program was differentially active in these cells. To get more insights into the differences at the molecular level between the Pre-HSPCs and Endo cells generated by 5TFs overexpression we decided to exploit the bioinformatic tool diffTF (Berest *et al*., 2019). Based on the ATAC-seq data and a set of predicted or validated TF binding sites, diffTF estimates which transcription factors are differentially active between two conditions (here: i5TFs Endo versus i5TFs Pre-HSPCs) (Fig.3C).

By applying diffTF to our data, we identified a number of transcription factors that appeared to be differentially active in i5TFs Endo versus i5TFs Pre-HSPCs (Fig. 3C). Some of the most active transcription factors in i5TFs Endo compared to i5TFs Pre-HSPCs were specific to VSM such as the SWI/SNF subunit *Smrc1*/*Smarcc1* and the Activator Protein-1 (AP-1) transcription factors *Fosl1*, *Fosl2*, *Jun, Junb, Jund* and *Batf* (Fig. 3C). Indeed, both families of proteins have known roles in the development and physiology of muscle cells, and the activity of these transcription factors was consistently found to characterize untreated eVSM cells in our diffTF analysis (Figure 3-figure supplement 2) (Andreucci *et al*., 2002; Zhou *et al*., 2009; Obier *et al*., 2016; Alonso *et al*., 2018). In contrast, one of the most active transcription factors in i5TFs Pre-HSPCs compared to i5TFs Endo was *Tal1* (Fig.3C). This unexpected finding was reinforced by the fact that *Tal1* was upregulated in i5TFs Pre-HSPCs but not in i5TFs Endo relative to i5TFs VSM (Fig.3D & 3E). *Lmo2* followed the same expression pattern (Fig. 3D & 3E). This finding suggests that the endogenous expression of *Tal1* and *Lmo2* may have a role in the generation of Pre-HSPCs following the overexpression of the 5TFs.

### 5TFs overexpression in hemangioblast cultures cannot generate Pre-HSPCs in the absence of a functional TAL1

Following the previous analysis, we hypothesized that the loss of *Tal1* would prevent the formation of Pre-HSPCs after the overexpression of the 5TFs. We focused on *Tal1* because it has been shown that the *Lmo2* ablation from mESCs allows the generation of Pre-HSPCs, albeit at lower frequency compared to *Lmo2^+/+^* mESCs, and *Lyl1* was not significantly upregulated in our datasets (Fig. 3E) (Stanulovic *et al*., 2017). To test our hypothesis, we decided to generate a *Tal1* knock-out in the i5TFs line genetic background. Using the CRISPR/CAS9 technology, we obtained two mutant lines (Fig. 4A & 4B). Differentiating these lines, we verified that the deletion in the *Tal1* gene prevented the formation of blood and endothelial cells, as only VSM cells were generated (Fig. 4C & 4D). This confirmed the absence of a functional TAL1 (Lancrin *et al*., 2009). Following the overexpression of the 5TFs in these cultures, we could only produce cells expressing the endothelial marker VE-CAD. No cells expressing CD41 were detected (Fig. 4C & 4D). Thus, we concluded that a functional TAL1 is required for the 5 other TFs to generate Pre-HSPCs from VSM.

**Figure 4:**
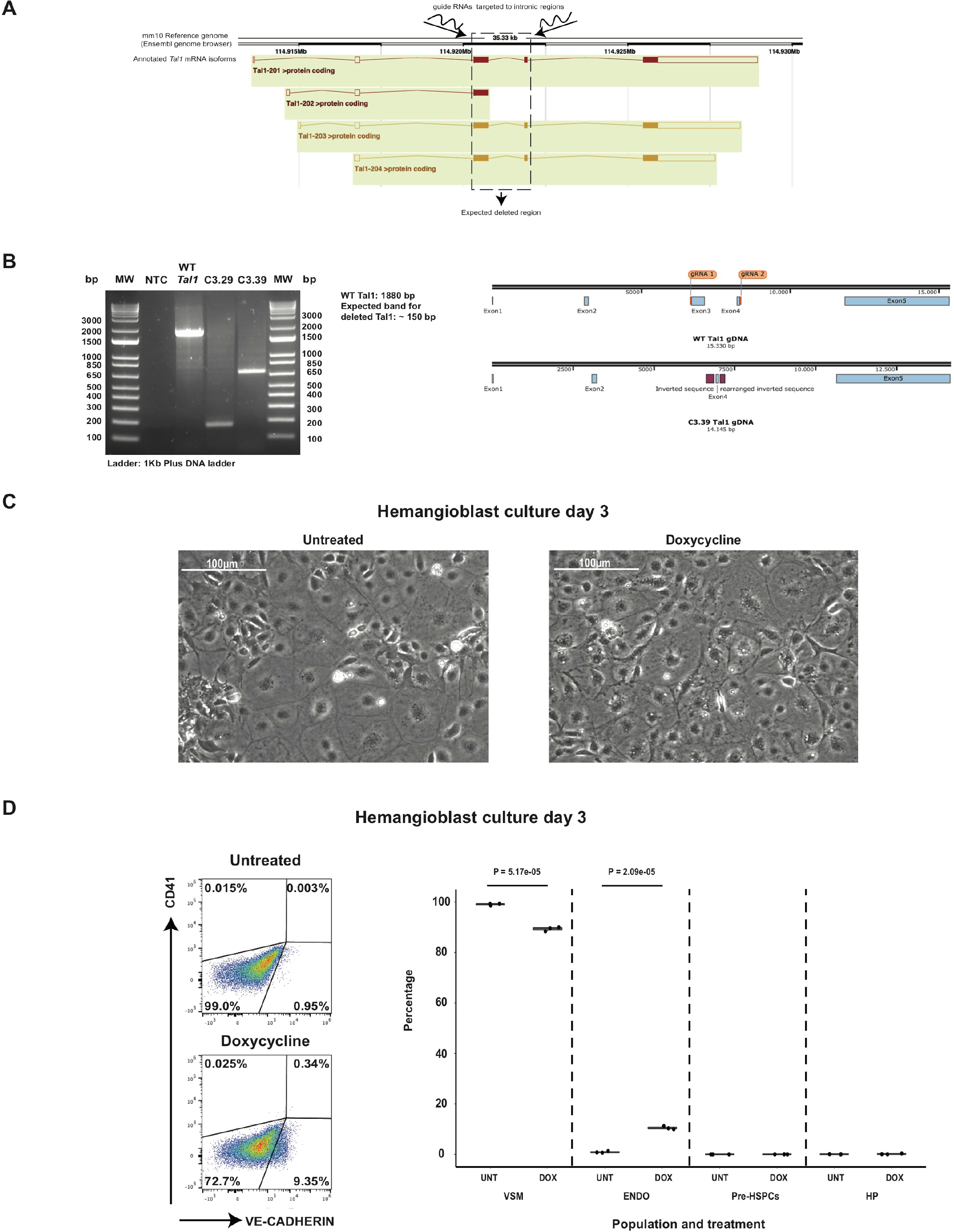
5TFs overexpression in hemangioblast cultures cannot generate Pre-HSPCs in the absence of a functional TAL1. A. CRISPR/Cas9 strategy used to disrupt the *Tal1* gene in the i5TFs mESC line (figure downloaded and modified from the Ensembl genome browser). B. Left: Electrophoresis showing the genotyping of the *Tal1* mutant mESCs clones used for experiments. Right: Scheme of the WT *Tal1* gene and the *Tal1* gene from mutant clone C3.39 reconstructed based on the sequencing results. C. Representative microscopy images of the indicated conditions at day 3 of hemangioblast culture. D. Representative flow cytometry analyses of the indicated conditions (left panels) and box plots showing the relative frequencies of VSM, Endo, Pre-HSPC and HP populations at day 3 of hemangioblast culture (n=3). Significance was determined by Analysis of Variance (ANOVA) test. Error bars correspond to standard deviations.

### Hierarchical clustering of differentially-expressed genes identified functional interactions between the 3TFs and the 5TFs on transcriptional regulation

Using the DESEq2 package (Love, Huber and Anders, 2014) we identified the differentially expressed genes (DEGs) between untreated controls and cells overexpressing the 8TFs, the 5TFs and the 3TFs (Supplementary file 10). To compare the effect of differential transcription factor overexpression we performed a hierarchical clustering analysis on the DEGs of the four sorted populations, dividing our dataset into 10 clusters containing genes with a similar expression pattern (Fig. 5A, Supplementary file 11). The clusters that were generated seem to reflect the relative contribution of the 5TFs and the 3TFs to the transcriptional changes induced by the 8TFs, and the acquisition of a Pre-HSPC-like identity. To exemplify this, cluster 1 contained mostly genes that were downregulated only in i8TFs Pre-HSPCs, suggesting that they were downregulated as a consequence of the combined action of the 5TFs with the 3TFs, while cluster 2 contained mostly genes that were clearly downregulated as a consequence of 5TF overexpression, with seemingly no contribution from the 3TFs. A similar analysis for each cluster can be found in Supplementary file 12. Such hierarchical clustering provided multiple pieces of evidence suggesting that a functional cooperation occurred among the 8TFs in regulating gene expression in i8TFs Pre-HSPCs. Indeed, we observed that a subset of genes was silenced (in cluster 1) and upregulated (in cluster 6) specifically when all 8TFs were overexpressed simultaneously, other genes were strongly silenced (in cluster 3) specifically when only the 5TFs but not the 8TFs were overexpressed, and a few genes (in clusters 6 and 10) appeared to be differentially regulated when the 3TFs were overexpressed alone or in combination with the other 5TFs. The hierarchical clustering analysis also highlighted transcriptional differences between the i5TFs Pre-HSPCs and the i8TFs Pre-HSPCs. This suggests that the expression of the 3TFs might be required for the proper transcriptional regulation of specific subsets of genes in Pre-HSPCs.

**Figure 5:**
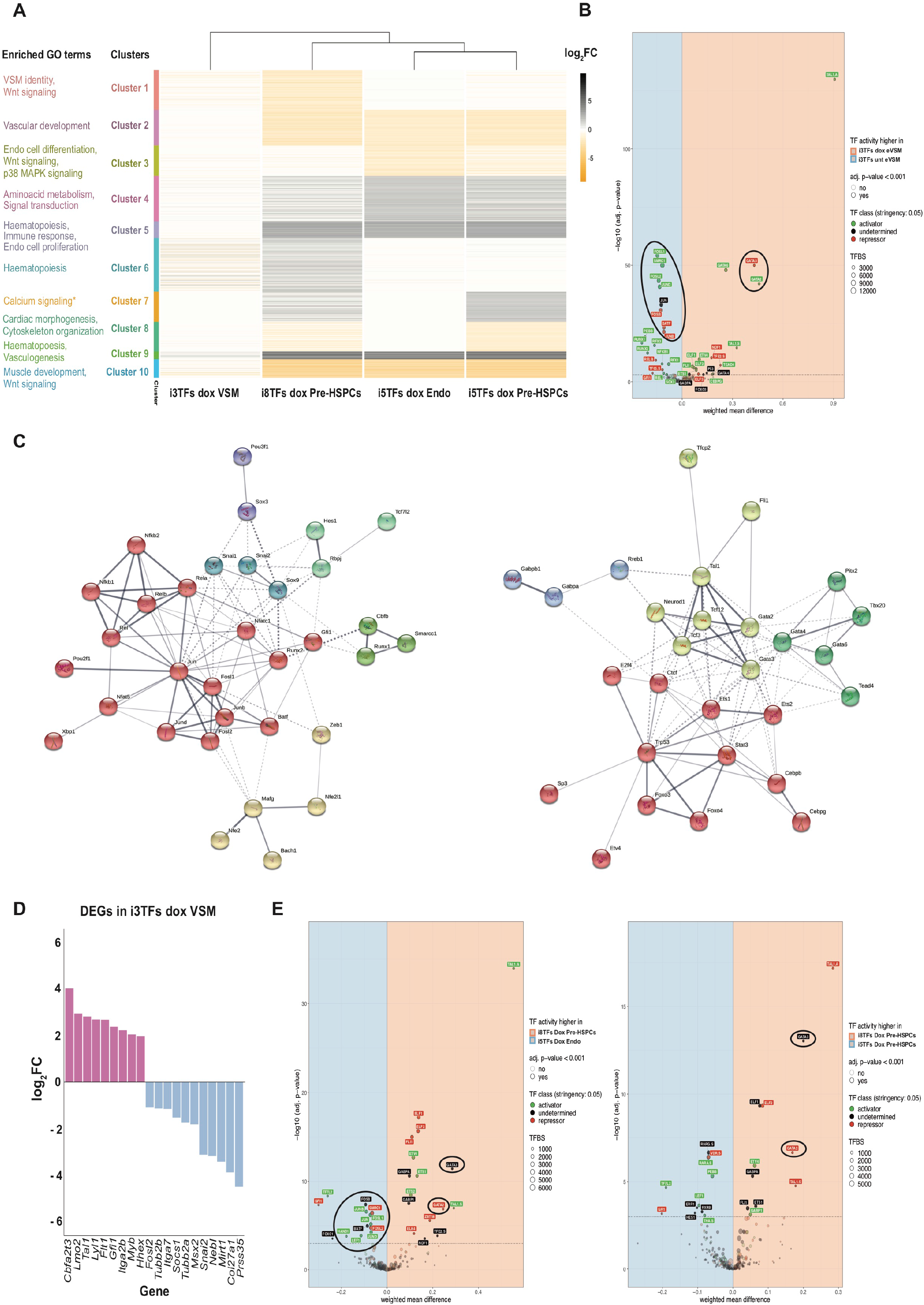
3TFs overexpression contributes to the silencing of the muscle transcriptional program and the activation of the hematopoietic one in eVSM. A. Heatmap showing the expression of genes from the clusters indicated in the figures. The most representative GO terms enriched in each cluster are shown in the left. * KEGG enrichment (no significant GO terms could be found for cluster 7). B. diffTF volcano plot comparing i3TFs untreated (unt) eVSM to i3TFs dox-treated (dox) eVSM. A detailed explanation of the plot can be found in Fig.3. TFs characteristic of VSM cells and hematopoietic TFs are circled. C. String interaction network analysis comparing i3TFs unt eVSM to i3TFs dox eVSM. D. Selection of differentially expressed genes (DEGs) comparing i3TFs unt eVSM to i3TFs dox eVSM. Hematopoietic genes are shown in pink, VSM-related genes are shown in blue. E. diffTF volcano plots comparing i8TFs Pre-HSPCs to i5TFs Pre-HSPCs (left panel) and i8TFs dox Pre-HSPCs to i5TFs dox Endo (right panel). A detailed explanation of the plots can be found in Fig.3. VSM and hematopoietic TFs are circled.

### 3TFs overexpression contributes to the silencing of the muscle transcriptional program and the activation of the hematopoietic one in eVSM

Next, we performed GO analysis on the genes from each cluster (Fig.5A, Figure 5-figure supplements 1 – 5, Supplementary files 13-21). The results of this analysis were consistent with the GO performed on the DARCs (Figure 3-figure supplement 1), and allowed us to identify a contribution of the 3TFs to the downregulation of non-hematopoietic transcriptional programs, particularly the muscle transcriptional program (clusters 1 and 10), and the upregulation of the hematopoietic one (clusters 6 and 9), as well as in the regulation of the Wnt, p38 Map kinase and BMP signaling pathways (clusters 3, 10). Indeed, we found cluster 1 to be enriched in GO terms related to functions that are important for muscle cells, such as extracellular matrix organization and muscle tissue development, as well as genes belonging to the Wnt signaling pathway (Fig. 5A, Figure 5-figure supplement 1A), while cluster 6 was predominantly enriched in GO terms related to hematopoiesis (Fig. 5A, Figure 5-figure supplement 1B). A detailed description of the GO terms enriched in each cluster can be found in Figure 5-figure supplements 1 – 5 and Supplementary files 13-21.

In agreement with the GO analysis, we found with our diffTF analysis combined with the use of STRING (Fig. 5B & 5C) that the smooth muscle specific TFs were more active in i3TFs unt eVSM compared to i3TFs dox eVSM. In contrast, several hematopoietic TFs were more active in the dox-treated conditions. Moreover, we found among the genes that were upregulated in i3TFs dox eVSM cells important hematopoietic regulators such as *Cbfa2t3*/*Eto2* (Schuh *et al*., 2005; Chagraoui *et al*., 2018), the *Tal1* target *Gfi1* (Wilson *et al*., 2010; Lancrin *et al*., 2012) and *Myb* (Mucenski *et al*., 1991) as well as the earliest murine hematopoietic marker *Itga2b*/Cd41 (Mikkola *et al*., 2003) (Fig. 5D), indicating that 3TFs overexpression alone was able to partially activate the hematopoietic transcriptional program. Likewise, among the downregulated genes we found ones with reported roles in vascular smooth muscle (*Mirt1* and *Msx2*) (Goupille *et al*., 2008; Li, Zhou and Huang, 2017), in myogenesis (*Snai2* and *Socs1*) (Diao, Wang and Wu, 2009; Soleimani *et al*., 2012), in directing hemangioblast cells towards the acquisition of a smooth muscle fate (*Fosl2*) (Obier *et al*., 2016) and genes that belong to classes that are generally important for the functionality of muscle cells, such as cytoskeleton remodeling proteins and extracellular matrix components, indicating that 3TFs overexpression alone was able to partially silence the muscle transcriptional program (Fig. 5D). Of note, nearly half of the DEGs in i3TFs dox eVSM cells were not in common with the DEGs in i8TFs Pre-HSPCs (Fig. 5-Figure Supplement 6), further supporting a functional interaction among the 8TFs that directs the transcriptional activity of these factors.

To gain further insights on the 3TFs role, we analyzed the diffTF results comparing the i5TFs line to the i8TFs one (Fig. 5E). When comparing the i8TFs Pre-HSPCs to the i5TFs Endo, the results appeared similar to the comparison between i5TFs Pre-HSPCs and the i5TFs Endo shown in Fig. 3C, with VSM TFs being the most active in i5TFs Endo and hematopoietic TFs being the most active in i8TFs Pre-HSPCs. When we compared the i8TFs Pre-HSPCs to the i5TFs Pre-HSPCs, however, we found that VSM TFs were no longer the most differentially active ones (Fig. 5E). Since the endogenous *Tal1* and *Lmo2* genes were upregulated in i5TFs Pre-HSPCs, this analysis provided orthogonal evidence suggesting that the upregulation of at least *Tal1* and *Lmo2* was sufficient to downregulate the VSM transcriptional program in the cells in which they were expressed.

### Rescue of *Tal1* expression during hemogenic endothelium culture allows the formation of Pre-HSPCs and HP from endothelium

Our previous analyses prompted us to assess whether *Tal1* played a role in the formation of Pre-HSPCs from the endothelium. The other studies that addressed this question did not find evidence to support a role for *Tal1* at this stage (Endoh *et al*., 2002; Schlaeger *et al*., 2005). To address this question, we developed a new system. The key was to be able to ensure the activity of *Tal1* in the mesoderm in order to produce endothelium and hemogenic endothelium, but to switch off its activity in these cell types in order to verify whether blood cell formation could progress. To make this new tool, we disabled the *Tal1* gene in the i3TFs line using the CRISPR/Cas9 method (Fig. 6A & 6B). As expected, no endothelial cells nor blood cells could be generated from these cell lines, confirming the absence of a functional TAL1 (Fig. 6B & 6D). To perform our experiments, we induced *Tal1*, *Lmo2* and *Lyl1* expression by addition of dox at day 2 of EB differentiation (Fig. 6C), as based on previous work (Liu *et al*., 2013), we expected this would be enough to produce FLK1^+^ mesodermal cells with hematopoietic potential. After sorting of FLK1^+^ cells, they were either plated with dox to maintain the expression of the 3TFs, or without to create a hemangioblast culture lacking the expression of a functional TAL1. Cells which were never exposed to dox (Unt Unt condition) were used as negative control. They were unable to produce any blood or endothelial cells but generated VSM as expected from cells lacking a functional TAL1. The constant treatment with dox (Dox Dox condition) led to a high frequency of HP after 2.75 days of culture. Endo and Pre-HSPCs could also be detected. In contrast, in the condition in which dox was only added at the EB stage (Dox Unt condition), the frequency of HP was about three times lower than in the Dox Dox condition (Fig. 6D). Hematopoiesis was therefore still ongoing after removal of dox albeit at a lower efficiency. At day 1.75 of culture *Tal1* expression in the Dox Unt condition was on average two times higher than the Unt Unt sample, but 256 times lower than in the Dox Dox condition (Fig. 7A). *Tal1* expression was therefore significantly reduced in Dox Unt cultures already 24 hours before the cytofluorimetric analysis, and virtually absent at day 2.75 (Fig. 7A).

**Figure 6:**
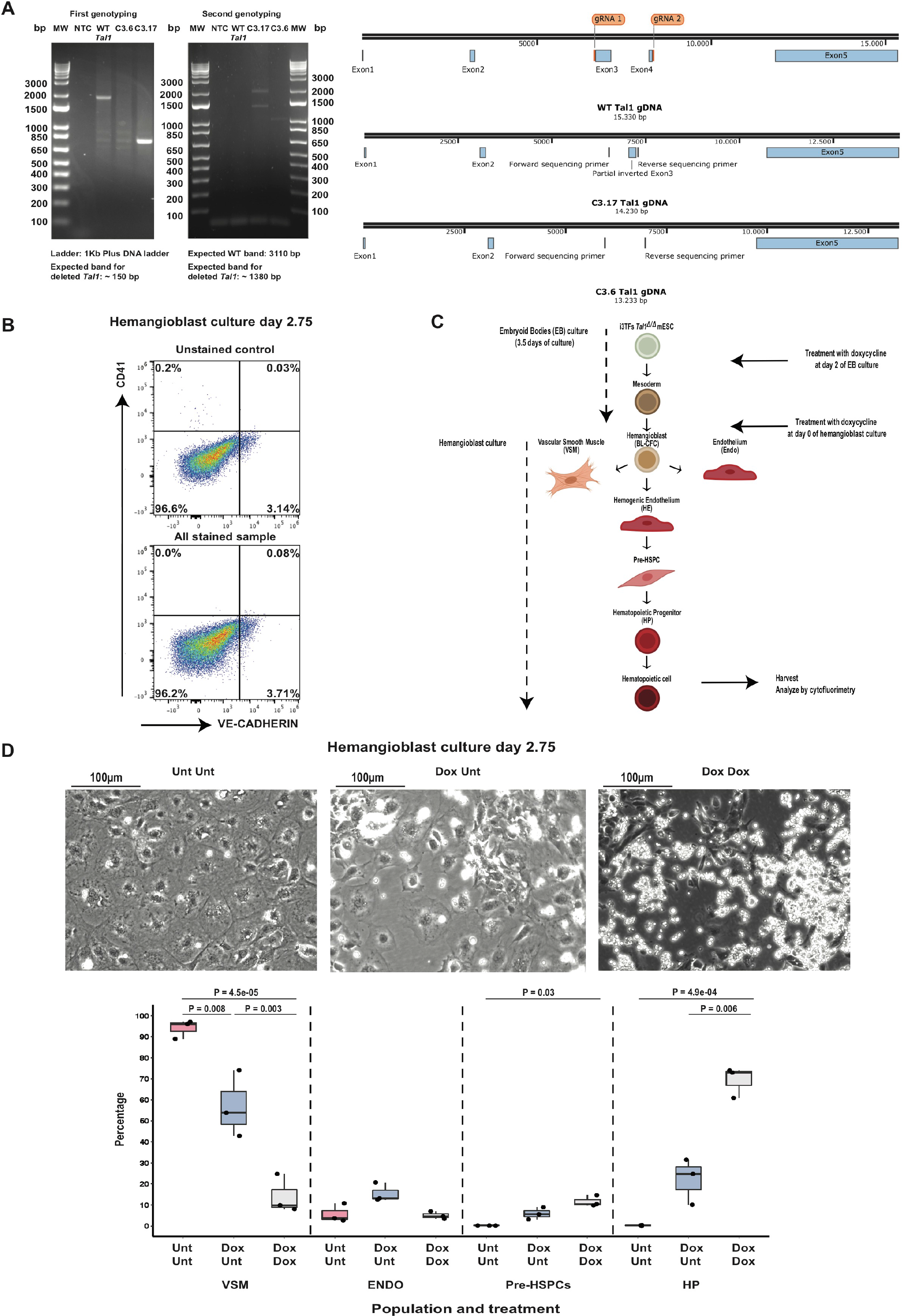
Characterization of the *Tal1 ^Δ/Δ^* inducible 3TFs embryonic stem cell line. A. Left: electrophoresis showing the genotyping of the *Tal1* mutant mESCs clones used for experiments. Right: scheme of the WT *Tal1* gene and the *Tal1* gene from mutant clones C3.17 (allele with the shorter deletion) and C3.6 reconstructed based on the sequencing results. B. Representative flow cytometry analyses of untreated day 1.75 hemangioblast cultures comparing the profile of unstained cells (top) and cultures stained for VE-CAD and CD41 (bottom). 3-4% of the cells in hemangioblast cultures were auto-fluorescent in the red (VE-CAD) channel. C. Experimental layout used to characterize the mutant line. D. Representative microscopic images of the indicated conditions. VSM cells appear as large wide-spread cell; Endo cells appear as elongated cells; HPs appear as round floating cells. Unt Unt = untreated; Dox Unt = doxycycline added at EB day 2; Dox Dox = doxycycline added at EB day 2 and at day 0 of hemangioblast culture. E. Box plots showing the relative frequencies of VSM, Endo, Pre-HSPC and HP populations of HE culture from the indicated conditions (n=3). Significance was determined by Analysis of Variance (ANOVA) test followed by Tukey HSD test for multiple test correction. Error bars correspond to standard deviations.

**Figure 7:**
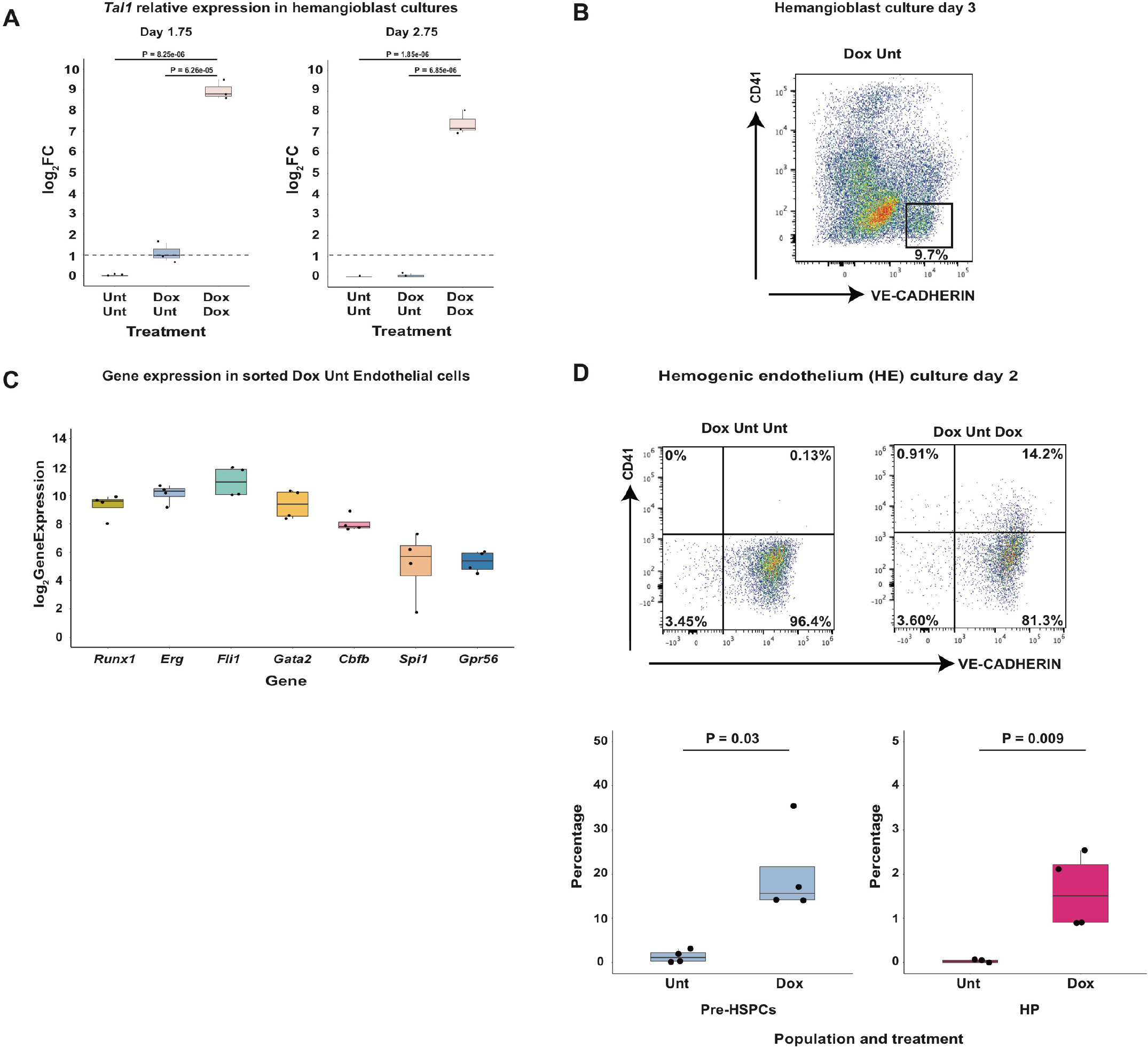
Rescue of *Tal1* expression during hemogenic endothelium culture allows the formation of Pre-HSPCs and HP from endothelium. A. qRT-PCR analysis showing the relative expression of the *Tal1* transgene transcript in the three highlighted conditions in day 1.75 (left) and day 2.75 (right) hemangioblast cultures. Significance was determined by Analysis of Variance (ANOVA) test on **Δ**Ct values. Error bars correspond to standard deviations. B. Representative flow cytometry analysis of hemangioblast cultures at day 3 highlighting the sorted endothelial cell population (VE-CAD^+^CD41^-^). C. RNA-seq (n=4) analysis showing the expression of the indicated genes in the FACS sorted endothelial cell population. D. Results of hemogenic endothelium cultures. Top: panels showing one representative flow cytometry analysis. Bottom: box plots summarizing four independent experiments. Significance was determined by Analysis of Variance (ANOVA) test. Error bars correspond to standard deviations.

We then continued with the hemogenic endothelium culture. We isolated VE-Cad^+^ CD41^-^ cells by FACS sorting at day 3 of hemangioblast culture. Parts of these cells were assessed by RNA-seq while the rest was used for the in vitro culture assay. We found that these cells strongly expressed *Runx1*, *Erg*, *Fli1*, *Gata2* and *Cbfb*, as well as additional hematopoietic genes such as *Gpr56*, which is required for HSPC generation (Kartalaei *et al*., 2015), and *Spi1/PU.1*, a regulator of HSPC differentiation (McKercher *et al*., 1996) (Fig. 7C, Supplementary file 22). The hemogenic endothelium culture was performed in absence or presence of dox. When the 3TFs were not induced, therefore in absence of *Tal1* expression, none or very few CD41^+^ cells were detected. In contrast, dox addition led to a consistent increase of CD41^+^ cell frequency. Most of these cells were VE-CAD^+^CD41^+^ (Pre-HSPCs), but more mature VE-CAD^-^CD41^+^ (HP) were also found. Thus, with this experiment we uncovered for the first time an unexpected requirement for *Tal1*in the generation of Pre-HSPCs during the process of EHT (Fig. 7D).

## DISCUSSION

The aim of our study was to elucidate the function of the hematopoietic regulators *Tal1*, *Lmo2* and *Lyl1* during the endothelial to hematopoietic transition (EHT). As their enforced expression in combination with the hematopoietic regulators *Runx1*, *Cbfb*, *Gata2*, *Erg* and *Fli1* was able to generate cells similar to Pre-HSPCs from mESC-derived hemangioblast cultures and mESC-derived VSM cells with very high efficiency (Bergiers *et al*., 2018), we addressed our question by investigating their contribution to Pre-HSPC production in these systems. We compared the effect of overexpressing just *Tal1*, *Lmo2* and *Lyl1* (3TFs) and just *Runx1*, *Cbfb*, *Gata2*, *Erg* and *Fli1* (5TFs) to the overexpression of all eight factors (8TFs) together, at the phenotypic and molecular level.

We found that overexpression of the 3TFs alone was able to trigger the activation of the hematopoietic transcriptional program and the downregulation of muscle-related genes in eVSM. This was further corroborated by our comparative analysis of the transcriptional changes induced by differential transcription factor overexpression. *Tal1* is essential for the hematopoietic commitment of mesodermal precursors (Dooley, Davidson and Zon, 2005; D’Souza, Elefanty and Keller, 2005; Lancrin *et al*., 2009), and multiple evidences showed that it has an important role in suppressing the alternative cardiac and paraxial lineage fates (Schoenebeck, Keegan and Yelon, 2007; Ismailoglu *et al*., 2008; Handel *et al*., 2012; Org *et al*., 2015; Chagraoui *et al*., 2018). However, a controversy still exists in the field on this matter, resulting from a sc-RNA-seq-based study on differentiating FLK1^+^ mesodermal cells from gastrulating embryos (Scialdone *et al*., 2016). Here, we have provided the first evidence that *Tal1*, in combination with its binding partner *Lmo2* and its homologous *Lyl1,* can promote the silencing of the VSM transcriptional program. This effect is likely to be mediated for the most part by *Tal1* in partnership with *Lmo2,* as their endogenous upregulation in i5TFs Pre-HSPCs appeared sufficient to down-modulate the activity of VSM transcriptional regulators according to our diffTF analysis. Our results are aligned with previous ones supporting a role for *Tal1* in the transcriptional repression of alternative fates, and suggest that *Tal1* is involved also in restricting VSM commitment of FLK1^+^ hemangioblast cells. Our main finding is that *Tal1* expression in endothelial cells is essential for Pre-HSPC generation. Instrumental for this was the use of eVSM cells as a platform for the comparative analysis of 5TFs and 8TFs overexpression, as it allowed a clear-cut readout without confounding effects deriving from a strong background expression of the assessed transcription factors. The different expression pattern of *Tal1* in i5TFs Endo and i5TFs Pre-HSPCs was indeed key to uncover the crucial role of this gene in the EHT process. We have previously shown by single-cell qPCR that a subset of VE-CAD^-^CD41^-^ cells in the hemangioblast culture at day 1 expressed low levels of *Tal1* (Bergiers *et al*., 2018). It is possible that the effect we observed here reflects a similar difference in the expression of *Tal1* in sorted FLK1^-^CD41^-^ eVSM cells, whereby cells that expressed this gene could acquire a Pre-HSPC phenotype upon 5TFs overexpression, whereas cells which did not acquired an Endo one. Another explanation derives from the multiple evidences that point to the existence of a regulatory loop among the 8TFs (Pimanda *et al*., 2007; Nicola K. Wilson *et al*., 2010; Ciau-Uitz *et al*., 2013; Mansour *et al*., 2014; Sanda and Leong, 2017), supporting a model in which the overexpression of the 5TFs determined the upregulation of *Tal1* which was sufficient in a subset of cells for the acquisition of a Pre-HSPC phenotype. Interestingly, this result could suggest that Pre-HSPC generation via overexpression of the 5TFs and the 8TFs occurred through an intermediate endothelial stage, mimicking the EHT. This is similar to what has been observed when hematopoietic precursors were generated from mouse fibroblasts via the enforced expression of *Gata2*, *Gfib*, *cFos* and *Etv6* (Pereira *et al*., 2013) or *Tal1*, *Lmo2*, *Runx1*, *Gata2* and *Erg* (Batta *et al*., 2014), and might indicate that passage through an endothelial stage is a *sine qua non* for the acquisition of a hematopoietic identity by non-hematopoietic cells.

Despite their prominent role in HSPC formation, we found that 5TFs expression at high levels in *Tal1****^Δ/Δ^*** endothelial cells was not sufficient for the generation of Pre-HSPCs from these cells, just like their overexpression was not sufficient to generate Pre-HSPCs in *Tal1****^Δ/Δ^*** hemangioblast cultures. This suggests that *Tal1* might be required in order for the 5TFs to exert their function in the EHT. Indeed, RUNX1, GATA2, ERG, FLI1, TAL1, LMO2 and LYL1 have been proposed to act as a multiprotein complex, and TAL1 is itself an obligate heterodimer the activity of which is largely modulated by protein-protein interactions (Hsu, Wadman and Baer, 1994; Wadman *et al*., 1997; Schuh *et al*., 2005; Goardon *et al*., 2006; Nicola K. Wilson *et al*., 2010; Stanulovic *et al*., 2017). It has been shown that TAL1 can recruit transcriptional regulators and chromatin remodeling complexes to its target genes through physical interactions, and similarly it can be directed to its target genes by interactions with other proteins (Schuh *et al*., 2005; Goardon *et al*., 2006; Stanulovic *et al*., 2017; Chagraoui *et al*., 2018). Thus, it is possible that TAL1 is required in hemogenic endothelial cells for the assembly of the multiprotein complex that contains the 5TFs, or to recruit transcriptional regulators that mediate essential functions of the 5TFs to this complex. Further experiments investigating the physical and functional interaction between TAL1 and the 5TFs would be of great value to further our understanding of the interplay among these important regulators of blood development.

From a translational viewpoint, our findings indicate that efforts to generate HSPCs *in vitro* or *ex vivo* should take in account the essential role of *Tal1* in Pre-HSPC generation, either favoring the use of starting cells which express this hematopoietic regulator, or exploiting alternative strategies to ensure its expression. Endothelial cells are particularly appealing for regenerative medicine purposes (Lis *et al*., 2017; Sugimura *et al*., 2017). By analyzing publicly available datasets of mouse scRNA-seq, we found that the frequency of adult endothelial cells expressing *Tal1* is very low compared to embryonic ones (Adamov *et al*., 2021). Based on our results, it appears therefore that any attempt to stimulate such cells to undergo EHT should include an increase in *Tal1* activity.

In conclusion, with our work we have uncovered an unexpected essential role for *Tal1* in the EHT. Additionally, we have shown that *Tal1*, together with *Lmo2* and *Lyl1*, concurs to activate the hematopoietic transcriptional program and suppress the muscle one in cells that transdifferentiate from VSM, providing ulterior evidence of the ability of *Tal1* to suppress alternative cell fates during hematopoietic development.

Through our bioinformatic analysis, we also identified the pathways and genes that were differentially expressed and differentially active in cells that acquired a Pre-HSPC phenotype upon 5TFs and 8TFs overexpression. Among these, there are likely to be some with regulatory roles in Pre-HSPCs. Our work therefore also constitutes a resource for the identification and investigation of potential novel regulators of Pre-HSPC formation and the EHT.

## MATERIAL AND METHODS

### EXPERIMENTAL MODEL AND SUBJECT DETAILS CELL LINES

#### Generation of inducible ESC lines

All doxycycline-inducible ESC lines were generated using the inducible cassette exchange method, the constructs and plasmids described previously (Iacovino *et al*., 2011; Vargel *et al*., 2016; Bergiers *et al*., 2018). The generation of the i8TFs mESC line is described in detail in Bergiers et al. For the generation of the i5TFs mESC line, HA-*Tal1*, FLAG-*Lyl1* and V5-*Lmo2* were successively excised from the p2lox-8TFs by classic cloning to generate the p2lox-5TFs used for the generation of the inducible ESC line. For the generation of the i3TFs mESC line, cmyc-*Fli1* and V5-*Erg* were successively excised from the intermediate p2lox construct containing Construct one and Construct two described in Bergiers et al. to generate the p2lox-3TFs used for the generation of the inducible ESC line. The functionality of the overexpression constructs of the chosen clones was corroborated by our RNA-seq and diffTF analyses (Fig.5D, Figure 3-figure supplement 2, Supplementary file 10).

#### Generation of cell lines with a deletion in the *Tal1* gene (i5TFs *Tal1^Δ/Δ^* mESCs, i3TFs *Tal1^Δ/Δ^* mESCs)

Two gRNAs targeting exon-flanking regions of introns 2 and 4 (relative to *Tal1*-201 mRNA) were used to generate a deletion in the *Tal1* gene (Fig. 4A & 4B, Fig. 6A). The intronic sequences were downloaded from the Ensemble genome browser v95 and used as input for the design of gRNAs. GRNAs were designed using the CRISPRko function of the sgRNA Design tool (now CRISPick) in the GPP Web Portal of the Broad Institute (https://portals.broadinstitute.org/gppx/crispick/public). The top candidate of the output list for each gene was selected for experiments. An extra G (in orange) before the sequence of the guide was added to the *Tal1* intron 2 guide to improve the efficiency of transcription from the U6 promoter. Sequences complementary to the overhangs created by digestion of the p133-pPB plasmid with BlpI and BstXI were added at both sides of the designed guides (sequences in blue). The guides were purchased from Sigma-Aldrich as sense and complementary antisense oligonucleotides.

*Tal1* intron 2: Sense 5’TTGGACGCACTGAAACCTGAAAAGGTTTAAGAGC3’; Antisense 5’TTAGCTCTTAAACCTTTTCAGGTTTCAGTGCGTCCAACAAG3’. *Tal1* intron 4: Sense 5’TTGGATGGTTCTAACCAGTGACAGTTTAAGAGC3’; Antisense 5’TTAGCTCTTAAACTGTCACTGGTTAGAACCATCCAACAA 3’.

To generate double-stranded oligonucleotides, 10µL of the sense and antisense oligonucleotides (100µM) were mixed with 80µL of 1.2X annealing buffer (10mM Tris pH 7.5-8.0, 60mM NaCl, 1mM EDTA), placed in a heating block at 95°C for 3’ and allowed to anneal at RT for at least 30’.

The guides were cloned individually by standard cloning into a p133-pPB plasmid containing an RNA Polymerase III-dependent U6 promoter, a modified gRNA stem loop and a BFP tag kindly provided by Jamie Hackett (EMBL Rome) (Figure 4-figure supplement 1).

The p133-pPB plasmids encoding for the *Tal1*-gRNAs were co-transfected with the pX458 plasmid containing a *Cas9* nuclease construct and a GFP reporter (pSpCas9-2A-GFP, Addgene ID: 48138), kindly provided by Jamie Hackett from EMBL Rome, to disrupt the *Tal1* gene in the i5TFs and i3TFs mESC lines.

MEFs were plated into gelatin-coated 6-well plates at a confluence of 0.2*10^6^ cells/well. The day before transfection, i5TFs and i3TFs mESCs were seeded on the MEFs at a confluence of 0.3*10^6^ cells/well in 3mL of DMEM-ES. Transient transfection was performed with polyethylenimine (PEI) based on the protocol of Longo and co-workers (Longo *et al*., 2013). Before transfection, the old medium was replaced with 2.7mL of fresh DMEM-ES. For each transfection, 9µg of PEI (kindly provided by the Genetic and viral engineering facility at EMBL-Rome) were diluted in DMEM in a final volume of 150µL. A total of 3µg of plasmid DNA per reaction were diluted in DMEM in a final volume of 150µL. The diluted PEI and DNA were mixed (3:1 ratio of PEI to DNA) and incubated at RT for 30’. The mixture was added to the cells dropwise. 1µg of each plasmid was used per reaction. Transfected cells were harvested after two days, the brightest BFP^+^GFP^+^ cells were FACS sorted (yields were between 5000 and 7000 cells) and plated in MEF-coated 10cm dishes in 10mL of DMEM-ES. 24 or 48 colonies were picked per cell line one week after sorting, dissociated by treatment with 30µL of TryplE Express for 3’ at 37°C and transferred into MEF-coated 96 well plates with 120µL of fresh DMEM-ES. The medium was changed the next morning to remove the TryplE. Growing clones were progressively transferred to MEF-coated 24-well and 6-well plates and frozen in 50% DMEM, 40% FBS and 10% DMSO, or frozen directly in a 96 well plate after splitting by adding 50µL of 2X freezing medium (80% FBS and 20% DMSO) to 40µL of dissociated cells and storing at -80°C. Cells from all clones were expanded in parallel on gelatin-coated plates for genotyping. The deletions in the *Tal1* gene were interrogated by PCR and sanger sequencing (Fig.4B, Fig.6A). The lack of functionality of the *Tal1* gene in the deleted clones was verified in the hemangioblast culture (Fig.4C & 4D, Fig.6D) (Lancrin et al., 2009). Two clones lacking *Tal1* functionality from both transfected mESC lines were selected for experiments (Fig.4B, Fig.6A).

#### Identification of cell lines

All cell lines used in this work were mESC lines. All A2lox.Cre-derived mESC lines were generated from the A2lox.Cre mESC line, which was a gift from Michael Kyba who produced it in his laboratory (Iacovino *et al*., 2011). All A2lox.Cre-derived mESC lines were generated in our laboratory as described in the section ‘Generation of inducible ESC lines’. All ESC lines had proper stem cell morphology and were able to give rise to blood, endothelial cells and vascular smooth muscle cells after in vitro differentiation (Fig.1C, Fig.2C).

#### Mouse embryonic stem cell (mESC) culture and embryoid bodies (EB) differentiation

All mESCs used in this work were maintained and expanded on a layer of MEFs in DMEM-ES medium composed of DMEM KO (supplemented with 1% Penicillin/Streptomycin, 1% L-glutamine, 1% non-essential amino acids), 15% FBS, 0.024% of LIF (1mg/mL) (produced by the protein expression facility at EMBL, Heidelberg) and 0.24% 50mM 2-Mercaptoethanol. All media were sterile-filtered before use in a Millipore stericup with a 0.22µm filter. Cells were incubated at 37°C with 5% CO 2 and 95% relative humidity.

Before plating of MEFs, culture dishes were gelatin coated by treatment with a solution of 0.1% gelatin in PBS for 20’ at room temperature (RT). MEFs were thawed at least 24hours before seeding of mESCs and plated at a confluence of 0.02*10^6^ cells/cm^2^ for expansion of mESC stocks and mESC line generation, and 0.017*10^6^ cells/cm^2^ for expansion of mESC prior to differentiation. MESCs were typically cultured in 6-well plates. Frozen cells were thawed in DMEM-ES and plated in one well in 4mL of DMEM-ES. Confluent cells were harvested and seeded at a density of 0.02-0.03*10^6^ cells/cm^2^ (0.2-0.3*10^6^ cells/well of a 6-well plate) in 2mL of medium for expansion, depending on the growth rate of the cells. The medium was changed daily to mESCs that were to be frozen as a stock. Cells were detached by treatment with TryplE Express (1mL/well of a 6-well plate) and 3-5’ of incubation at 37°C.

For differentiation into EBs, mESCs were subjected to two successive passages on gelatin-coated dishes for the removal of MEFs. Confluent cells at the center of the well were harvested following a 2’ RT incubation with TryplE Express pre-heated to 37°C, to minimize the detachment and carryover of MEFs. 1.8*10^6^ cells/dish were plated into gelatin-coated 10cm dishes in 10mL of DMDM-ES. Cells were harvested after 24hours and plated for the second gelatin passage in 10mL of IMDM (Iscove’s Modified Dulbecco’s Medium) -ES medium composed of IMDM (supplemented with 1% Penicilliin/Streptomycin, 1% L-glutamine), 15% FBS, 0.024% of LIF (1mg/mL) and 0.24% 50mM 2-Mercaptoethanol. Cells were harvested after 24hours and plated into standard 90mm petri dishes at a confluence of 0.3*10^6^ cells/dish, in 10mL of EB medium composed of IMDM (supplemented with 1% Penicilliin/Streptomycin and 1% L-glutamine), 15% FBS, 0.6% Transferrin, 0.03% monothioglycerol (MTG) and 50µg/mL ascorbic acid. Cells were kept in culture for 3-3.5 days and then harvested for BL-CFC sorting. Cultures were checked at day 2 and transferred to new petri dishes if the EBs were attaching to the bottom of the dishes. For the *Tal1^Δ/Δ^* i3TFs cell line, doxycycline was added to day 2 EBs to a final concentration of 1µg/mL.

#### Method details

##### Flow cytometry and cell sorting

Staining was performed as described in Oatley *et al*., 2020, or as described in the “Hemogenic endothelium (HE)” section. Cells from EBs, hemangioblast, hemogenic endothelium and OP9/OP9-DL1 cultures were stained with different combinations of antibodies. Dyes 7AAD (Invitrogen, A1310) or Sytox Blue (Invitrogen, S34857) were used to exclude dead cells. FACS analysis was performed using a FACSCanto (Becton Dickinson) and an Attune NxT Flow Cytometer (Thermo Fisher Scientific). Cell sorting was performed using the FACSAria (Becton Dickinson) or by using magnetic sorting (MACS MicroBead Technology, Miltenyi Biotec) and anti-APC MicroBeads (Miltenyi Biotech). Data were later analyzed using FlowJo v10.1r5 (Tree Star, Inc.).

##### Hemangioblast culture

For hemangioblast cultures, MACS sorted FLK1^+^ BL-CFCs were plated on gelatin-coated plates at a confluence of 0.0105*10^6^ cells/cm^2^ (freshly sorted cells) and 0.0168*10^6^ cells/cm^2^ (frozen cells) in a hemangioblast medium containing IMDM (supplemented with 1% Penicillin/Streptomycin and 1% L-glutamine), 10% FBS, 0.6% human transferrin, 0.3% MTG, 50µg/mL ascorbic acid, 0.05% VEGF (10µg/mL) and 0.1% IL-6 (10µg/mL). 15% D4T supernatant was also added to the medium in experiments performed with the i3TFs, i5TFs, i8TFs mESCs. This is the supernatant of endothelial D4T cells cultured in IMDM media with 10% FBS and 30mg of endothelial growth supplement. BL-CFCs were cultured in hemangioblast medium for up to 3 days, and were harvested at different time-points for downstream analyses. Doxycycline was added to the cells to a final concentration of 1µg/mL.

##### Hemogenic endothelium (HE) culture

FLK1^-^CD41^-^ VSM cells from day 1 hemangioblast cultures and VE-CAD^+^CD41^-^ endothelial cells from day 3 hemangioblast cultures were FACS-sorted and plated in gelatin-coated dishes at a density of 0.0095*10^6^ cells/cm^2^. Cells were cultured up to 2 days in a hematopoiesis-promoting HE medium containing IMDM (supplemented with 1% Penicillin/Streptomycin and 1% L-glutamine), 10% FBS, 1% L-glutamine, 0.6% human transferrin, 0.3% MTG, 50µg/mL ascorbic acid, 0.024% of LIF (1mg/mL), 0.5% SCF (10µg/mL), 0.1% oncostatin M (10µg/mL) and 0.01% FGF (10µg/mL). Doxycycline was added to the FLK1^-^CD41^-^ VSM cells to a final concentration of 10µg/mL. Doxycycline was added to the VE-CAD^+^CD41^-^ endothelial cells to a final concentration of 1µg/mL.

For FACS and flow cytometry analysis, cells were stained with the antibodies shown in table 3.

**Table 1:** Reagents and commercial kits.

| Product or resource | Supplier | Reference number |
| --- | --- | --- |
| **Antibodies** |  |  |
| Anti-APC MicroBeads | Miltenyi Biotec | 130-090-855 |
| Anti-CD144 eFluor-660, eBioBV13 | eBioscience | 50-1441-82 |
| Anti-Mouse CD309 APC, Avas12a1 | eBioscience | 17-5821-81 |
| Anti-Mouse CD41 PE, MWReg30 | eBioscience | 12-0411-81 |
| **Chemicals** |  |  |
| 100X SYBR Green I | Invitrogen | S7563 |
| 2-mercaptoethanol | GIBCO | 31350-010 |
| Ascorbic acid | Sigma-Aldrich | A4544 |
| Betaine | Sigma-Aldrich | 61962 |
| CompBead Anti-Rat and Anti-Hamster Ig $\kappa$ /Negative Control Compensation Particles | BD Biosciences | 552845 |
| D4T supernatant | EMBL-Rome | N/A |
| DMEM KnockOut (KO) | GIBCO | 10829-018 |
| Doxycycline | Sigma-Aldrich | D9891 |
| DTT | Invitrogen | 18064-014 |
| EPO | R and D | 959-ME-010 |
| Fetal bovine plasma-derived serum platelet poor (PDS) | Antech | N/A |
| Fetal bovine serum (FBS) | GIBCO | 10270106 |
| Gelatin | BDH | 440454B |
| GM-CSF | R and D | 425 ML |
| Human FGF basic | R and D | 233-FB-025 |
| IGEPAL CA 630 | Sigma-Aldrich | I8896 |
| KAPA SYBR FAST qPCR Master Mix (2X) Kit | KAPABIOSYSTEMS | KR0389_S |
| KAPA2G Robust HotStart ReadyMix PCR Kit | KAPABIOSYSTEMS | KR0381_S |
| IL-11 | R and D | 418 ML |
| IL-3 | R and D | 403 ML |
| IL-6 | R and D | 406 ML |
| IMDM | Lonza | BE12-726F |
| L-glutamine | GIBCO | 25030-024 |
| LIF | EMBL Heidelberg | N/A |
| LightCycler 480 SYBR Green I Master | Roche | 04707516001 |
| Magnesium chloride ($\text{MgCl}_2$) | Sigma-Aldrich | M8266 |
| MCSF | R and D | 416 ML-010 |
| MEM Non-Essential Amino Acids Solution | GIBCO | 11140-035 |
| Methylcellulose | VWR | 9004-67-5 |
| MinElute PCR Purification Kit | QIAGEN | 28004 |
| Monothioglycerol (MTG) | Sigma-Aldrich | M6145 |
| NEBNext High-Fidelity 2X PCR Master Mix | New England Biolabs | M0541 |
| Nextera Index Kit | Illumina | FC-121-1011 |
| Oncostatin M | R and D | 495-MO |
| Penicillin-streptomycin | GIBCO | 15140-122 |
| PFMH-II | GIBCO | 12040-077 |
| Recombinant RNase inhibitor | Clontech | 2313A |
| RNeasy Micro Kit | QIAGEN | 74004 |
| RNeasy Plus Mini Kit | QIAGEN | 74134 |
| SCF | R and D | 455-MC |
| SPRISelect | Beckman Coulter | B23318 |
| Superscript II reverse transcriptase | Invitrogen | 18064-014 |
| TD (2X reaction buffer from Nextera kit) | Illumina | FC-121-1030 |
| TDE (Nextera Tn5 transposase from Nextera kit) | Illumina | FC-121-1030 |
| TPO | R and D | 488-TO-005 |
| Transferrin | Roche (Italy) | 10652202001 |
| Triton X-100 | Sigma-Aldrich | T9284 |
| TrypLE express | GIBCO | 12605-036 |
| TWEEN 20 | Sigma-Aldrich | P1379 |
| VEGF | R and D | 293-VE |

**Table 2:**
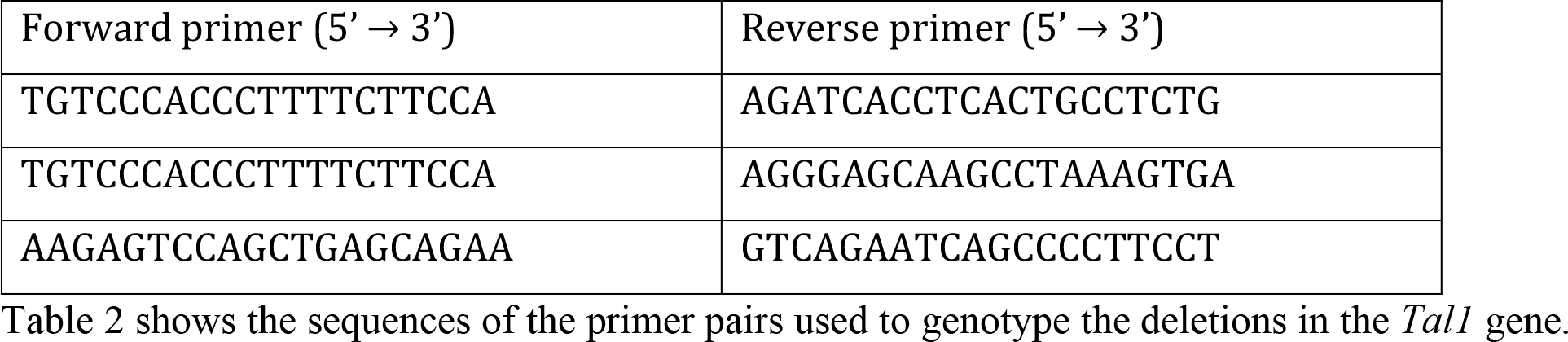
Primer pairs for genotyping of Tal1.

**Table 3:** Antibodies used for flow cytometry analysis and FACS sorting.

| Antigen | Fluorochrome | Working dilution | Clone | Reference |
| --- | --- | --- | --- | --- |
| CD309<br>(FLK1) | APC | 1:300 | Avas12a1 | eBioscience;<br>17-5821-81 |
| CD144<br>(VE-CADHERIN) | eFluor-660 | 1:200 | eBioBV13 | eBioscience;<br>50-1441-82 |
| CD41a<br>(CD41) | PE | 1:400 | MWReg30 | eBioscience; 12-0411-82 |
| CD117<br>(C-KIT) | BV421 | 1:200 | 2B8 | BD Biosciences;<br>562609 |

**Table 4:**
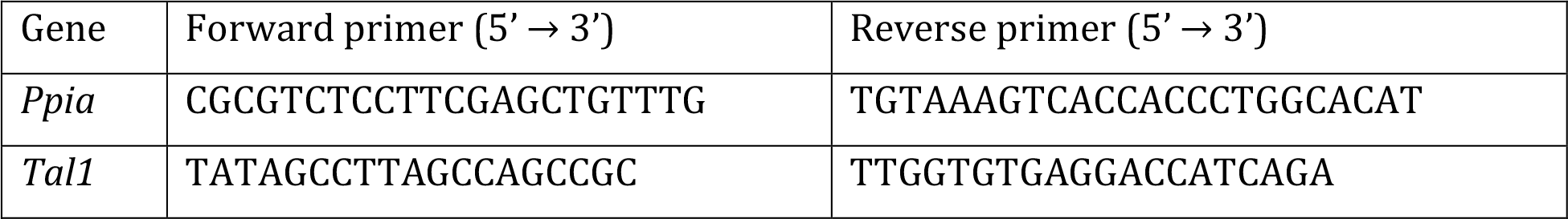
Primer pairs used for qPCR.

Unstained, single stained and fluorescence minus one (FMO) controls were prepared and analyzed during each experiment. Compensation beads (BD Biosciences) were used for single-stained controls. Cells heat-shocked at 60°C for 5’ were used for the 7-AAD single-stained control. 0.1 or 0.05*10^6^ cells were typically used for controls. 5000 cells were used for controls in experiments where the cell numbers were limiting (analysis of the hemogenic endothelial cultures from sorted VE-CAD^+^CD41^-^ cells). Separate sets of Unstained and FMO controls were typically prepared for untreated and dox-treated conditions. When the cell numbers were limiting (analysis of the hemogenic endothelial cultures from sorted VE-CAD^+^CD41^-^ cells) cells from both conditions were pooled together for the controls. 0.1*10^6^ cells were typically stained for flow cytometry analysis, and at least 30.000 events per sample were acquired. When cell numbers were limiting, all the remaining cells after aliquoting of the controls were stained, and the maximum number of events possible per sample was acquired.

Cells were stained in a final volume of 50 or 60µl of staining mix (FACS buffer with antibodies). For FACS sorting, all the cells remaining after aliquoting of the controls were stained, and the maximum possible number of cells was sorted. Cells for sorting were stained in 200µl of staining mix per 10^6^ cells.

For staining, 2X or 3X (depending on the number of antibodies used for staining) antibody solutions were prepared, and equal volumes of each were mixed to obtain a final 1X staining mix. FMO controls were stained by adding directly 25µl of 2X antibody solution or 20µl of 3X antibody solutions with an equal volume of FACS buffer to the tubes. For staining, cells were resuspended in staining mix and incubated for 10’ at RT, in the dark shaking at 500 RPM. Cells were washed after staining with 1mL of FACS buffer to remove unbound antibodies, and resuspended in FACS buffer with 7-AAD (100X) diluted 1:100 (volume/volume ratio). Unstained controls were resuspended in 50 or 60µl of FACS buffer.

Compensation beads were stained by adding 1 drop of negative beads (CompBead Negative negative Control), 1 drop of positive beads (CompBead Anti-Rat/Hamster Ig, k beads) and 50 or 60µl of staining mix into a tube and incubating in the dark at RT for 5’ or in ice for 20’.

##### Genomic DNA (gDNA) extraction

gDNA extraction for genotyping was performed on snap-frozen culture-derived cells using the DNeasy Blood & Tissue Kit and the QIAmp DNA Micro Kit from Qiagen, according to manufacturer’s instructions. DNA samples were stored at -20°C.

##### Genotyping of cell lines

PCRs for the genotyping of cell lins were performed using the KAPA2G Robust HotStart ReadyMix according to manufacturer’s instructions using an annealing temperature of 60°C. The primers used for each gene were described previously. Reactions were set-up mixing 6.25µl of 2X KAPA2G master mix, 0.625µl of 10µM forward primer, 0.625µl of 10µM reverse primer, 100ng of gDNA and nuclease-free H2O to a final volume of 12.5µl.

##### RNA extraction

RNA extraction was performed on snap-frozen culture-derived cells using the RNeasy Plus Mini Kit and RNeasy Micro Kit according to manufacturer’s instructions, based on the amount of starting material. Prior to RNA extraction, cells were homogenized using a syringe and 20-gauge needle or the QIAshredder homogenizer. The RNA was stored at -80°C.

##### Reverse transcription and cDNA production

cDNA production was performed using the RevertAid H Minus RT cDNA Synthesis kit (ThermoFisher Scientific) according to manufacturer’s instructions, starting from 200ng of RNA. The reaction was set-up mixing 1µl of random hexamer primer, 4µl of 5X reaction buffer, 1µl of RiboLock Rnase Inhibitor (20U/µl), 2µl of dNTP mix 10mM, 1µl of RevertAid H Minus M-MuLV Reverse Transcriptase (200 U/µL) and 200ng of RNA with nuclease-free H2O for molecular biology in a final volume of 20µl. The reaction was incubated in a thermocycler for 5’ at 25°C, 60’ at 42°C and 70°C for 5’. The cDNA was stored at -20°C.

##### Quantitative PCR

qPCR reactions were performed with a 7500 Real-Time PCR System from Applied Biosystems, using the KAPA SYBR FAST ROX low qPCR Master Mix (2X) Kit. The reactions were set-up mixing 5µl of KAPA SYBR FAST qPCR Master Mix (2X) with 0.4µl of 10µM forward primer, 0.4µl of 10µM reverse primer, 2µl of cDNA and 3.2µl nuclease-free H2O in a final volume of 10µl.

Reactions were cycled as follows: hold at 50°C for 2’, enzyme activation at 95°C for 10’, 40 cycles of exponential amplification at 95°C for 15’’, 60°C for 1’ (signal acquisition was performed at this stage), 1 cycle of amplification for melting curve analysis at 95°C for 30’’, 60°C for 1’, 95°C for 30’’ (signal acquisition was performed between the two last stages), and 60°C for 15’’.

The mouse house-keeping gene *Ppia* (*Peptidylprolyl Isomerase A*) was used as an internal control for normalization in all qPCR reactions.

The primer pairs used for qPCR are listed in Table 5

##### Assay for Transposase-Accessible Chromatin using sequencing (ATAC-seq)

ATAC-seq was performed based on the protocol described by Buenrostro and coworkers (Buenrostro *et al*., 2015) 5000 cells were sorted directly into 1.5mL Eppendorf tubes in 50µL of PBS 10% FBS. Cells were centrifuged at 500 x g for 5’ at 4°C, the supernatant was carefully removed and cells were gently resuspended in 100µL of cold (pre-cooled at 4°C) lysis buffer (10mM Tris-HCl, pH 7.4, 10mM NaCl, 3mM MgCl2, 0.1% NP40/IGEPAL CA-630, 0.1% Tween-20). Cells were immediately centrifuged at 500 x g for 10’ at 4°C. The supernatant was removed and cells were washed once with 50µL of cold 1 x PBS and centrifuged at 500 x g for 5’ at 4°C. The supernatant was carefully removed, the nuclei were resuspended in 50µL of transposition mix composed of 25µL of TD (2X reaction buffer), 2.5µL of TDE (Nextera Tn5 Transposase) and 22.5µL of molecular grade nuclease-free H2O, and incubated at 37°C for 30’. The transposed DNA was purified using the Qiagen MinElute PCR Purification Kit following manufacturer’s instructions and eluted in 10µL of Elution Buffer (10mM Tris buffer, pH 8). The purified product was frozen and stored at -20°C for later amplification of the transposed regions. The entire volume (10µL) of eluted DNA was used for PCR amplification with 2.5µL of Custom Nextera PCR primer 1 (containing barcodes), 2.5µL of Custom Nextera PCR primer 2 (containing barcodes), 25µL of NEBNext High-Fidelity 2X PCR Master Mix and 10µL of molecular-grade nuclease-free H2O for a final 50µL reaction. The reaction was thermal cycled as follows: 1 cycle at 72°C for 5’, 1 cycle at 98°C for 30’’, 5 cycles of exponential amplification at 98°C for 10’’, 63°C for 30’’ and 72°C for 1’. 5µL of the reaction were used for qPCR to determine the remaining number of amplification cycles for each sample. For qPCR, 5µL of amplified DNA were mixed with 0.5µL of PCR Primer Cocktail, 0.09µL of 100X SYBR green I, 5µL of NEBNext High-Fidelity 2X PCR Master Mix and 4.41µL of molecular-grade nuclease-free H2O for a final 15µL reaction. The qPCR was cycled as follows: 1 cycle at 98°C for 30’’, 20 cycles of exponential amplification at 98°C for 10’’, 63°C for 30’’ and 72°C for 1’. The additional number of cycles needed per sample was calculated by plotting the linear Rn value versus the number of cycles and determining the cycle number that corresponds to 1/3 (one third) of the maximum fluorescence intensity.

The remaining 45µL of the PCR reaction were amplified for the number of cycles calculated with the qPCR (1 cycle at 72°C for 5’, 1 cycle at 98°C for 30’’, N cycles of exponential amplification at 98°C for 10’’, 63°C for 30’’ and 72°C for 1’) and stored at -20°C. The amplification products were purified using SPRI beads (Beckman Coulter) according to the manufacturer’s instructions, using a bead-to-DNA ratio of 1.1X, and eluted in 22µL of TE Buffer (TrisHCl 10mM, pH 8).

Purified samples were analyzed by capillary electrophoresis on a 2100 Bioanalyzer using the Agilent High Sensitivity DNA Kit to verify the quality of the samples and quantify the samples before pooling. Samples were mixed equimolarly into two pools with 10 samples each. The final libraries were analyzed by capillary electrophoresis on the Bioanalyzer to verify the quality of the libraries and quantify them before sequencing. The libraries were paired-end sequenced with the Illumina technology on a NextSeq500 (2 x 75bp read length, mid-output) at the Genomics Core Facility at EMBL Heidelberg.

##### RNA sequencing (RNA-seq)

RNA-Seq was performed according to the SmartSeq2 protocol described by Picelli and coworkers (Picelli *et al*., 2014). 25 cells were FACS sorted directly into 0.2mL safe-lock microtubes in 4µL of a freshly-prepared solution containing 2µL of lysis buffer (1µL of RNase Inhibitor 2U/µL with 19µL of a 0.2% (vol/vol) Triton X-100 solution), 1µL of oligo-dT30VN primer 10µM and 1µL of dNTP mix 10mM. The tubes were quickly vortexed, spun down, immediately snap-frozen in dry ice and stored at -80°C for later processing. The steps of cDNA conversion, pre-amplification, purification with magnetic beads and quality control were performed at the Genomics Core Facility at EMBL Heidelberg or in house. The quality of the cDNA samples was assessed by capillary electrophoresis on a 2100 Bioanalyzer using the Agilent High Sensitivity DNA Kit. The sequencing library preparation and sequencing were performed at the Genomics Core Facility at EMBL Heidelberg. Libraries containing 24 samples each were paired-end sequenced with the Illumina technology on a NextSeq500 (2 x 75bp read length, high-output).

#### Quantification and statistical analysis

##### Flow cytometry analysis

All flow cytometry experiments were independently repeated three times. The box plots were generated using the R software. One-way ANOVA (Analysis of variance) was performed on the cell frequencies. Error bars correspond to standard deviations.

##### ATAC-seq data analysis

For the processing of the ATAC-seq sequencing data an in-house *Snakemake* pipeline was used for all processing steps starting from the raw fastq files from sequencing. The details of the data processing are described in the “ATAC-seq processing” paragraph of the methods section of the manuscript of Berest, Arnold and co-workers (Berest, Arnold *et al*., 2019).

The processed BAM files and BED files were used for pair-wise comparisons of chromatin accessibility between conditions using the *R*/Bioconductor package DiffBind version 2.8.0 (Stark and Brown, 2011). For differential analysis the default setting was used. The scoring function used for dba.count was DBA_SCORE_RPKM and the data were normalized on the library size.

##### Gene Ontology (GO) analysis on ATAC-seq data

GO analysis was performed using the GREAT (Genomic Regions Enrichment Annotation Tool) online tool from Stanford University (McLean *et al*., 2010). The BED files obtained from the DiffBind analysis, containing the regions of chromatin differentially accessible between doxycycline-treated samples and untreated controls, were sorted to keep only regions with a log2FC > | 2 | and a false-discovery rate (FDR) adjusted p-value < 0.05 (supplementary file 1). These regions were used as input for the analysis, using the following parameters: species assembly Mouse GRCm38 (UCSC mm10, Dec.2011), background regions whole genome, association rules setting Basal plus extension (Proximal: 5kb upstream, 1kb downstream, plus Distal: up to 1000.0 kb), Include curated regulatory domains selected.

##### diffTF analysis on ATAC-Seq data

diffTF version 1.6 was used with RNA-seq integration, Hocomocov version 11, and cluster.largeAnalysis.json. The following options were set in config.json: “maxCoresPerRule”: 20, “nPermutations”: 0, “nBootstraps”: 1000, “nCGBins”: 10, “TFs”: “all”. The documentation can be found at https://difftf.readthedocs.io/en/v1.6/. The details of the data processing for the diffTF analysis of the ATAC-Seq sequencing data are described in the manuscript of Berest, Arnold and co-workers (Berest, Arnold *et al*., 2019).

##### String analysis on diffTF results

For String analysis, we used the output of the diffTF analysis corresponding to a TF class stringency of 0.001 and adj. p-value < 0.05. We filtered out the TFs that were not expressed according to out RNA-seq analysis on the corresponding cells. For the generation of the interaction networks we used the web tool STRING at https://string-db.org/, and displayed only the TFs that were connected to at least another TFs in the network. For the network generation, we selected the following basic settings: meaning of network edges “confidence”, active interaction sources “Textmining, Experiments, Databases, Co-expression, Neighborhood, Gene Fusion, Co-occurrence”, minimum required interaction score “medium confidence (0.400)”, max number of interactions to show “1^st^ shell -none/query proteins only, 2^nd^ shell -none”. We selected the following advanced settings: network display mode “interactive svg”, display simplifications “hide disconnected nodes in the network”. For the clustering method we selected “MCL clustering” with an inflation parameter of 3.

##### RNA-seq data analysis

The processing of the RNA-seq data was performed using the EMBL Galaxy Server (https://galaxy.embl.de/) (Afgan *et al*., 2018). FastQC was used to perform quality control on the raw sequencing data. Removal of adaptor sequences (trimming) from paired-end reads was performed with Trim Galore!. FastQC was run again to perform quality control on the trimmed reads. Reads were aligned to the reference mouse genome GRCm38 (UCSC mm10) using the RNA STAR (Spliced Transcripts Alignment to a Reference) Aligner (Dobin *et al*., 2013). Filter SAM or BAM was used to filter the BAM files using SAMtools to remove poor quality alignments and retain one alignment per read (Skip alignments with any of these flag bits set: the alignment of this read is not primary, the read fails platform/vendor quality checks, supplementary alignment) (Li *et al*., 2009). bamCoverage (deepTools) was used to generate a coverage bigWig file (bin size in bases: 10, Scaling/Normalization method: 1x) (Ramírez *et al*., 2016). featureCounts was used to count how many reads have mapped to genes (measure gene expression from BAM files) (Liao, Smyth and Shi, 2014). CollectRNASeqMetrics (Picard tools) was used to collect metrics about the alignment of RNA to various functional classes of loci in the genome (http://broadinstitute.github.io/picard/). CollectInsertSizeMetrics (Picard tools) was used to plot the distribution of insert sizes (http://broadinstitute.github.io/picard/).

Differential expression analysis was performed using the DESeq2 package with the R software (version 3.5.1, http://www.R-project.org.) along with R studio (version 0.99.879) (Team, 2008; Love, Huber and Anders, 2014).

##### Hierarchical clustering of RNA-seq data and GO analysis on the clusters

The hierarchical clustering analysis of the differentially expressed genes (DEGs) of the four dox-treated populations, and the GO analysis on the 10 identified clusters was performed using the R software (Team, 2008). All DEGs identified by DESeq2 from the pairwise gene-expression comparisons between untreated controls and dox-treated samples were merged into one omni-comprehensive matrix. Genes in the matrix that were not identified as differentially expressed in one of the conditions were assigned a log2FC value of 0 replacing the “NA” (not available) value. A distance matrix was calculated using the Manhattan distance, and used to perform the clustering into 10 groups using Ward’s minimum variance method to measure the dissimilarity between two clusters of observations (Murtagh and Legendre, 2014). The clustering was performed with the R stats function hclust, and a heatmap was generated with the R function pheatmap (Kolde, 2019).

GO analysis and KEGG enrichment were performed using the R package clusterProfiler (Durinck *et al*., 2009; Yu *et al*., 2012). Emaps for the visualization of GO and KEGG analyses were created using the enrich plot package (Yu, 2021).

#### Data and software availability

##### Software

All software is freely or commercially available and is listed in the Methods description and product reference table.

## Acknowledgments

We thank Cora Chadick and Christopher Hall (EMBL Rome FACS Facility, Italy) for cell sorting; Vladimir Benes (EMBL, Genomics Core Facility, Heidelberg, Germany) for next-generation sequencing; Valentina Carlini (EMBL, Italy) and Cristina Policarpi (EMBL, Italy) for advice on CRISPR/CAS9 gene editing; Ana Boskovic (EMBL, Italy), Katharina Koch (EMBL, Italy) for critical reading of the manuscript; Kim Dale (University of Dundee, Scotland), Jamie Hackett (EMBL, Italy) and Karima Kissa (University of Montpellier, France) for fruitful scientific discussions. The European Molecular Biology Laboratory supported this work.

## Author Contributions

Yasmin Natalia Serina Secanechia, Conceptualization, Formal analysis, Investigation, Supervision, Visualization, Writing—original draft, Writing—review and editing; Isabelle Bergiers, Conceptualization, Formal analysis, Writing—review and editing, Supervision; Matt Rogon, Formal analysis, Visualization, Writing—review and editing; Christian Arnold, Formal analysis, Visualization, Writing—review and editing; Nicolas Descostes, Formal analysis, Visualization,Writing—review and editing; Stephanie Le, Investigation, Writing—review and editing; Natalia Lopez Anguita, Investigation, Writing—review and editing; Kerstin Ganter, Investigation, Writing—review and editing; Chrysi Kapsali, Investigation, Writing—review and editing; Lea Bouilleau, Investigation, Writing—review and editing; Aaron Gut, Investigation, Writing—review and editing; Auguste Uzuotaite, Investigation, Writing—review and editing; Ayshan Aliyeva, Investigation, Writing—review and editing; Judith Zaugg, Supervision, Investigation, Writing— review and editing; Christophe Lancrin, Conceptualization, Formal analysis, Supervision, Investigation, Visualization, Methodology, Writing—original draft, Project administration, Writing—review and editing.

