## Supplementary material for "Identifying a novel role for the master regulator *Tal1* in the Endothelial to Hematopoietic Transition": Figure 3 - figure supplements 1 -2

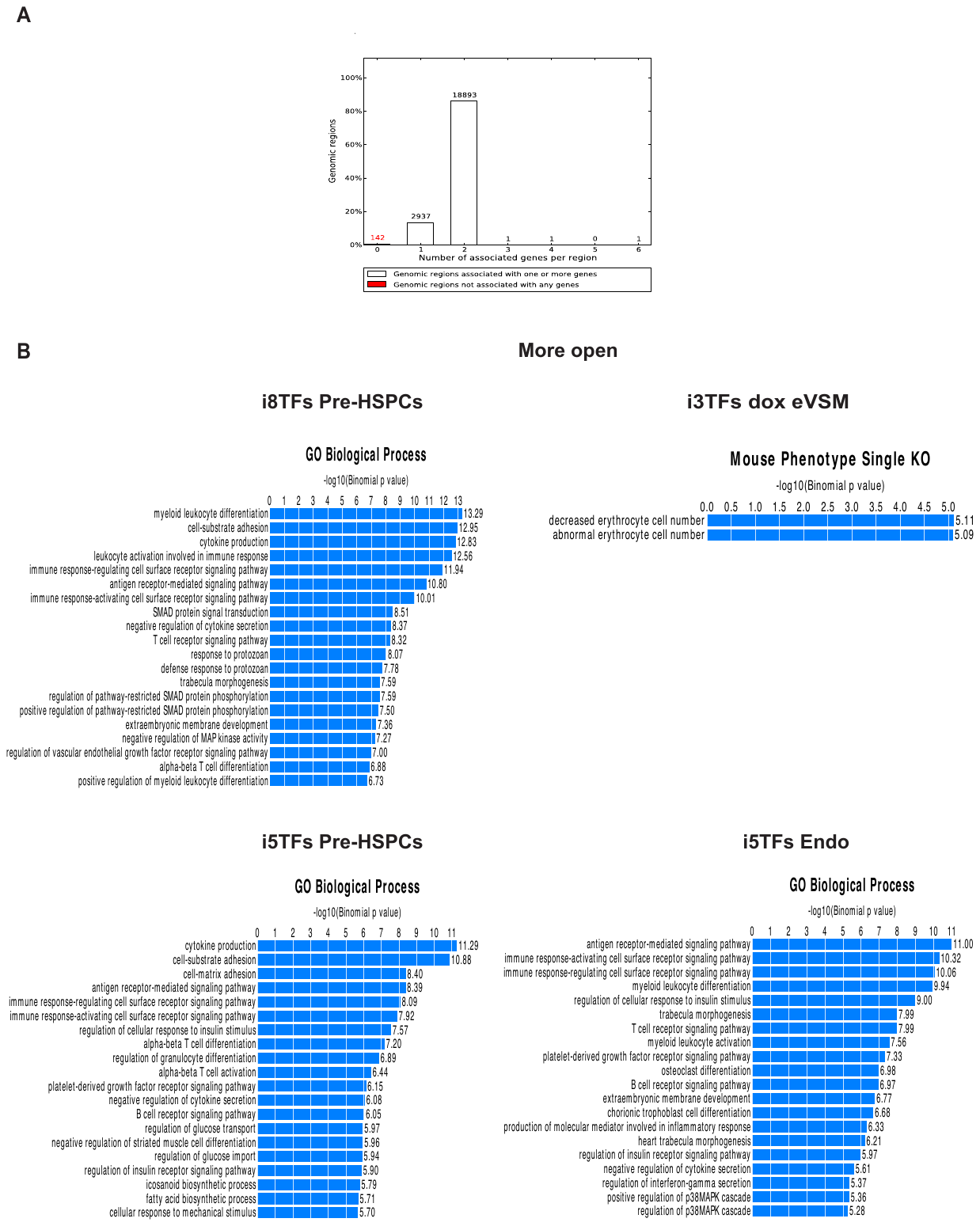

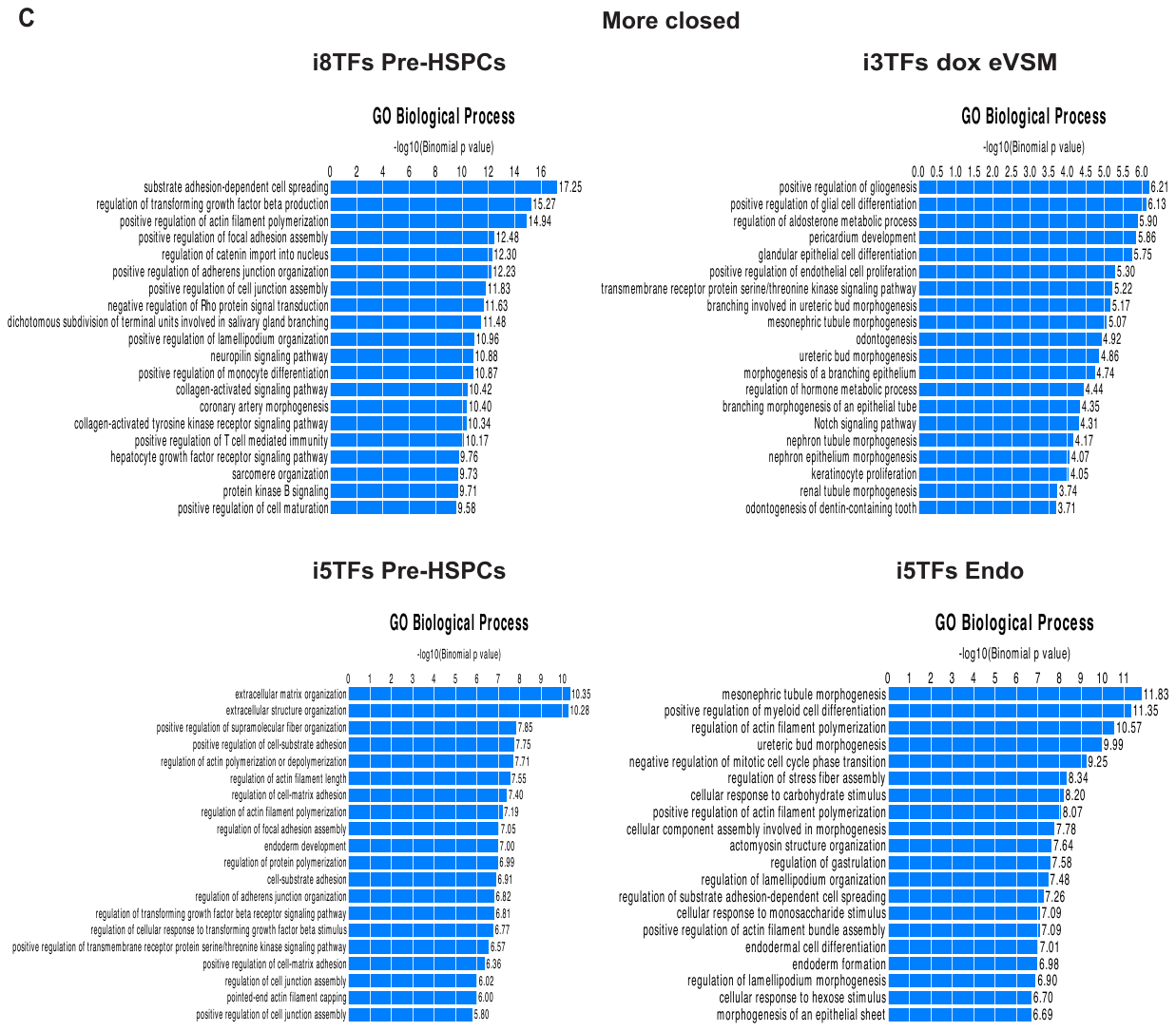

**Figure 3-figure supplement 1: GO terms enriched at differentially accessible regions of chromatin**

**A.** Bar-plots showing the number of genes associated with the differentially accessible regions of chromatin identified by DiffBind. **B.** Bar-plots showing the GO terms enriched at regions of chromatin more open in dox-treated cells compared to untreated controls. **C.** Bar-plots showing the GO terms enriched at regions of chromatin more closed in dox-treated cells compared to untreated controls.

**Figure 3-figure supplement 2: Volcano-plots showing the transcription factors identified as differentially active in dox-treated cells and untreated controls**

Volcano-plots showing the transcription factors (TFs) identified by diffTF (Berest, Arnold *et al*., 2019) as differentially active in untreated cells (blue quadrant) and dox-treated cells (red quadrant). TFs classified as activators are labelled in green, TFs classified as repressors are labelled in red, TFs that couldn’t be classified as either are labelled in black. 5% of TFs were classified as activators or repressors (TF class stringency: 0.05) based on the Pearson correlation index. The x axis (weighted mean difference) shows the difference in TF activity between the untreated (unt) and dox-treated (dox) conditions. The y-axis displays the significance of the TFs. The significance threshold is indicated with a dotted line (FDR adjusted P-value < 0.05). Transcription-factor binding sites (TFBS) are indicated as a dot. The size of each dot is proportional to the number of genomic TFBS identified for each TF. VSM-specific transcription factors are highlighted.
