## Supplementary material for "Identifying a novel role for the master regulator *Tal1* in the Endothelial to Hematopoietic Transition": Figure 4 - figure supplement 1

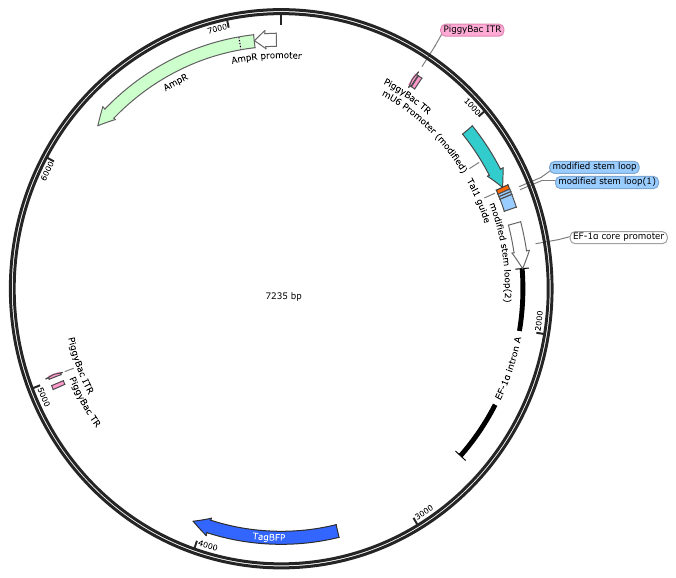

**Figure 4-figure supplement 1: Scheme of the p133-pPB plasmid encoding for the Tal1 gRNA used for the disruption of the *Tal1* gene**
