## Supplementary material for "Identifying a novel role for the master regulator *Tal1* in the Endothelial to Hematopoietic Transition": Figure 5 - figure supplement 1-6

**Figure 5-figure supplement 1: GO terms enriched in clusters one and six**

Enrichment maps (EMaps) showing the GO terms enriched in clusters 1 (**A**) and 6 (**B**) identified by the hierarchical clustering of DEGs in i8TFs Pre-HSPCs, i5TFs Pre-HSPCs, i5TFs Endo and i3TFs dox eVSM. Each node (circle) represents a GO term. The size of the nodes is proportional to the number of genes belonging to each GO term, and the color represents the significance. The edges (lines) connect terms with shared genes, so GO terms with mutually overlapping gene sets tend to cluster together.

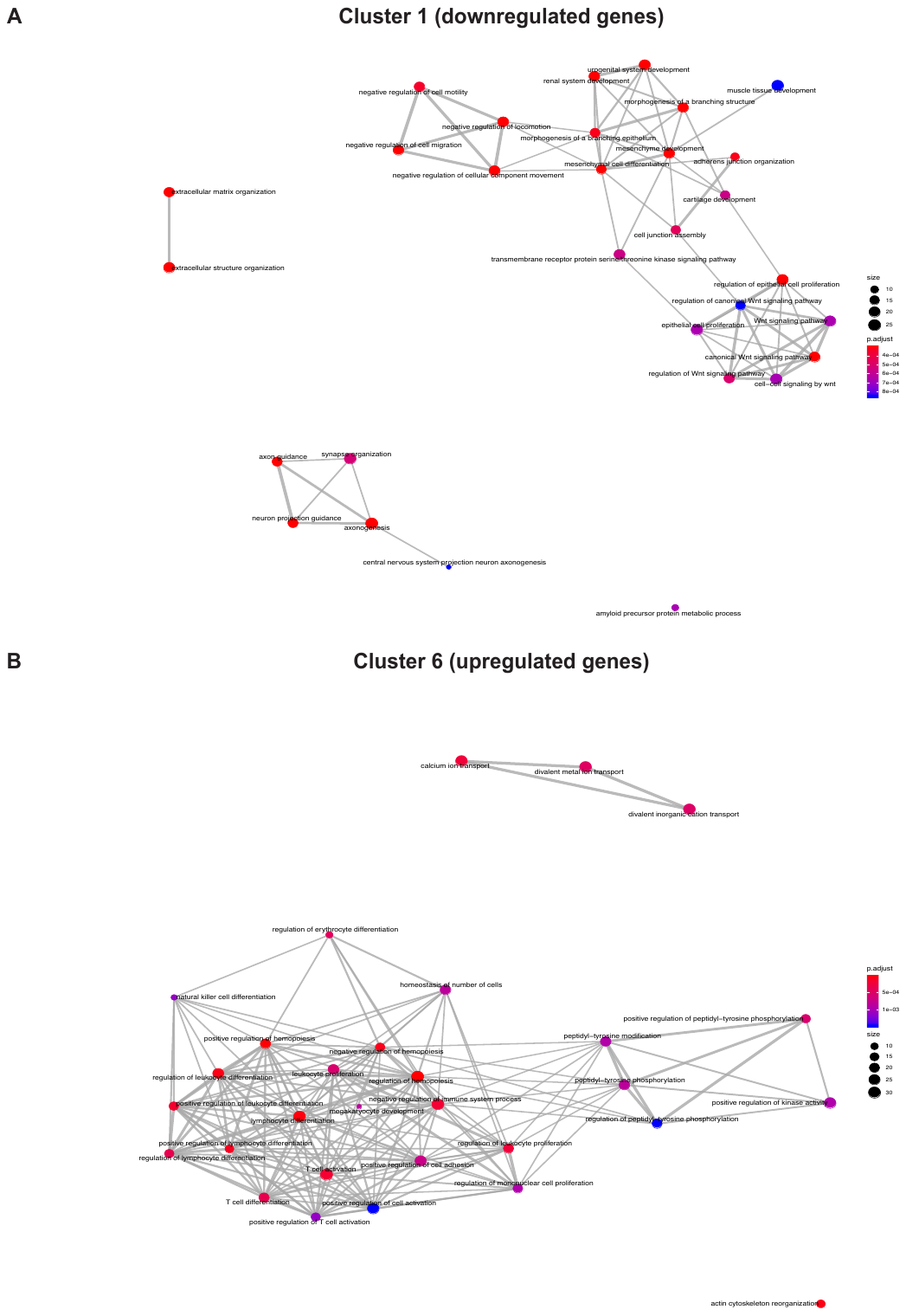

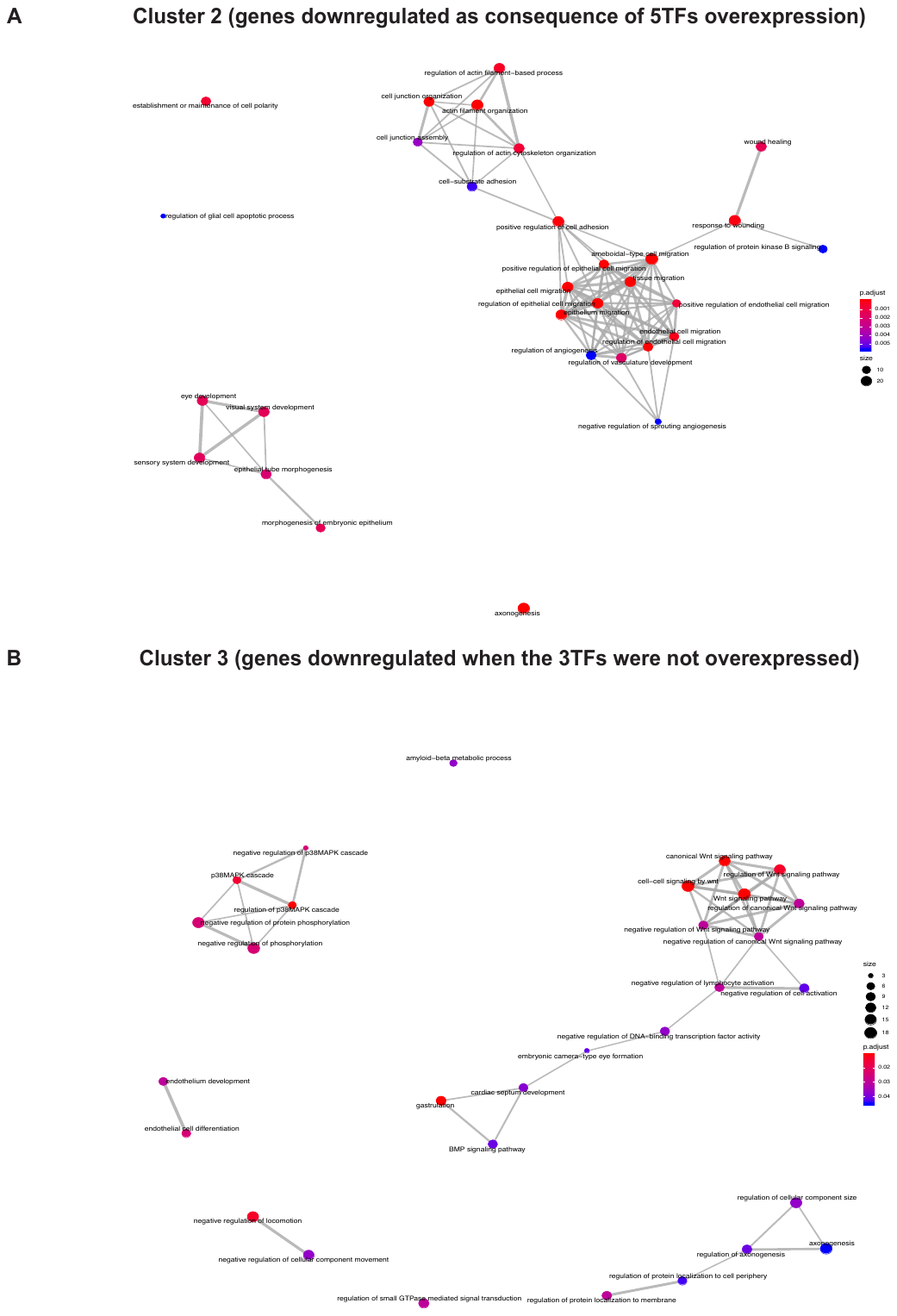
**Figure 5-figure supplement 2: GO terms enriched in clusters two and three**

Enrichment maps (EMaps) showing the GO terms enriched in clusters 2 (**A**) and 3 (**B**) identified by the hierarchical clustering of DEGs in i8TFs Pre-HSPCs, i5TFs Pre-HSPCs, i5TFs Endo and i3TFs dox eVSM. Each node (circle) represents a GO term. The size of the nodes is proportional to the number of genes belonging to each GO term, and the color represents the significance. The edges (lines) connect terms with shared genes, so GO terms with mutually overlapping gene sets tend to cluster together.

**Figure 5-figure supplement 3: GO terms enriched in clusters four and five**

Enrichment maps (EMaps) showing the GO terms enriched in clusters 4 (**A**) and 5 (**B**) identified by the hierarchical clustering of DEGs in i8TFs Pre-HSPCs, i5TFs Pre-HSPCs, i5TFs Endo and i3TFs dox eVSM. Each node (circle) represents a GO term. The size of the nodes is proportional to the number of genes belonging to each GO term, and the color represents the significance. The edges (lines) connect terms with shared genes, so GO terms with mutually overlapping gene sets tend to cluster together.

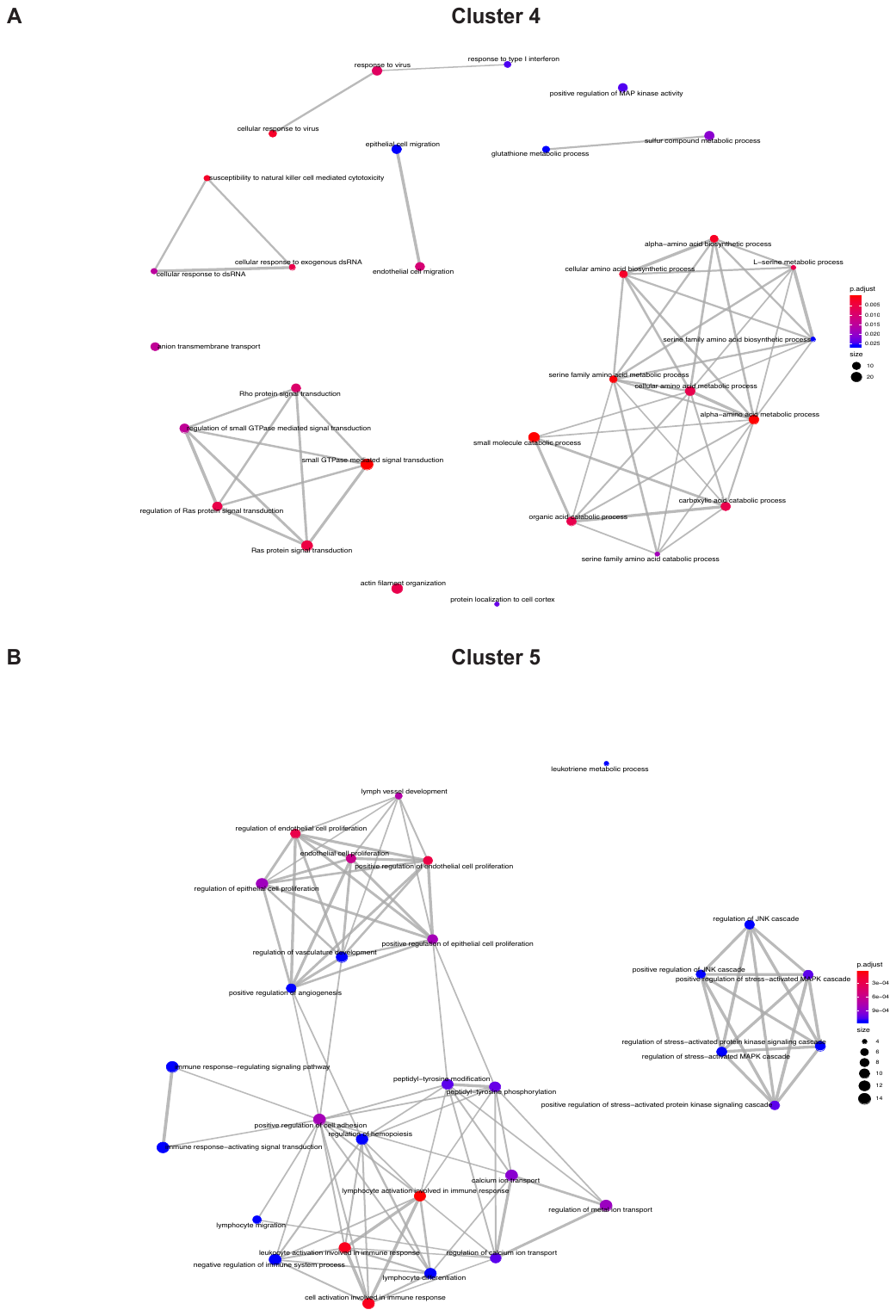

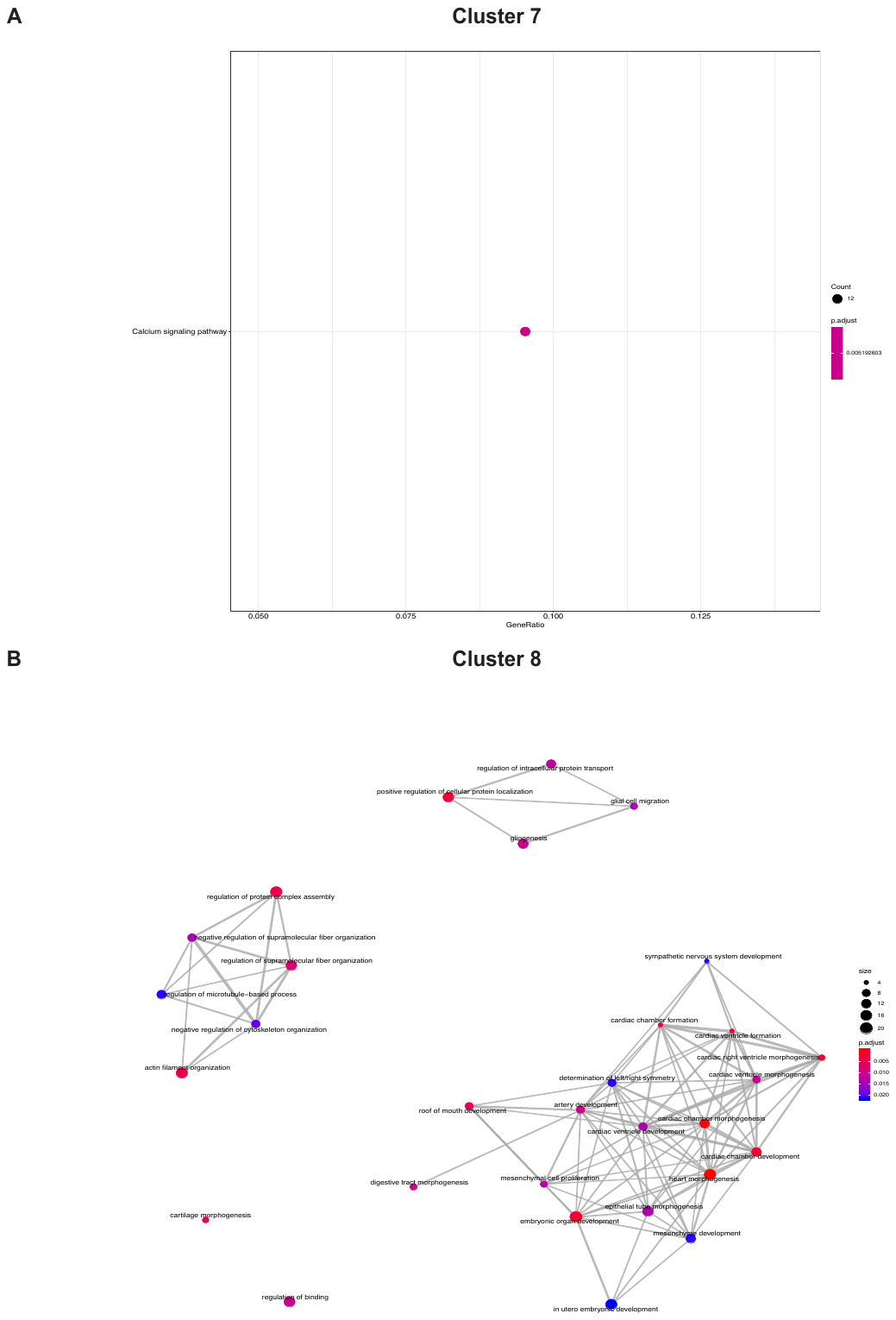

**Figure 5-figure supplement 4: GO terms enriched in clusters seven and eight**

Dotplot and Enrichment map (EMap) showing the GO terms enriched in clusters 7 (**A**) and 8 (**B**) identified by the hierarchical clustering of DEGs in i8TFs Pre-HSPCs, i5TFs Pre-HSPCs, i5TFs Endo and i3TFs dox eVSM. Each node (circle) represents a GO term. The size of the nodes is proportional to the number of genes belonging to each GO term, and the color represents the significance. The edges (lines) connect terms with shared genes, so GO terms with mutually overlapping gene sets tend to cluster together.

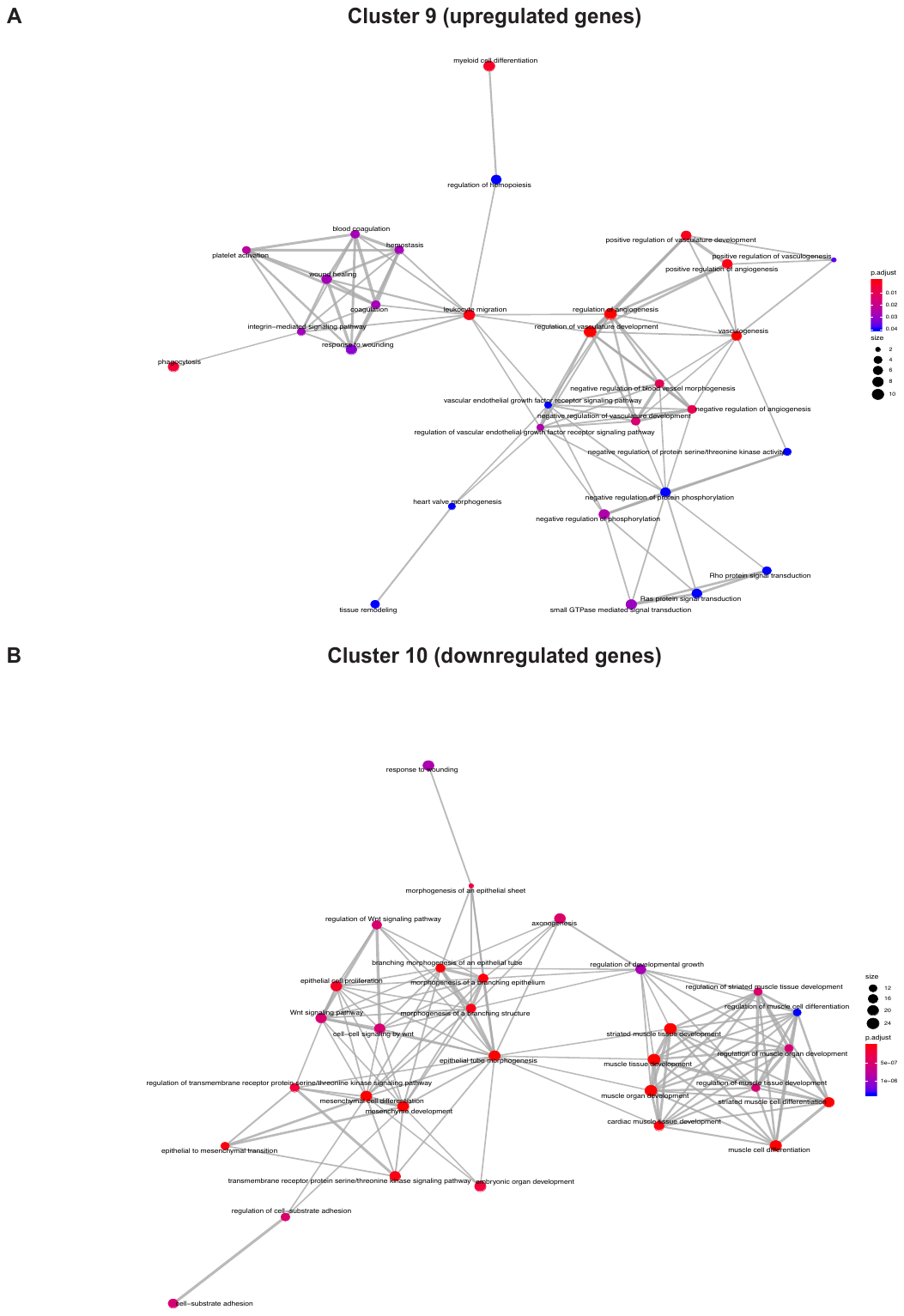

**Figure 5-figure supplement 5: GO terms enriched in clusters seven and eight**

Enrichment maps (EMaps) showing the GO terms enriched in clusters 9 (**A**) and 10 (**B**) identified by the hierarchical clustering of DEGs in i8TFs Pre-HSPCs, i5TFs Pre-HSPCs, i5TFs Endo and i3TFs dox eVSM. Each node (circle) represents a GO term. The size of the nodes is proportional to the number of genes belonging to each GO term, and the color represents the significance. The edges (lines) connect terms with shared genes, so GO terms with mutually overlapping gene sets tend to cluster together.

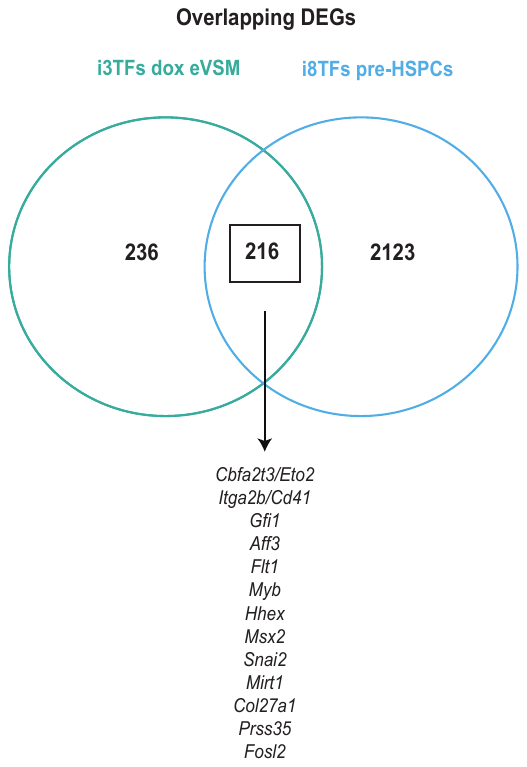

**Figure 5-figure supplement 6: DEG overlap between i3TFs dox eVSM and i8TFs Pre-HSPCs**

Venn diagrams comparing the DEGs in i3TFs dox eVSM and i8TFs Pre-HSPCs. A subset of differentially expressed hematopoietic and muscle-related genes in common between the two conditions is shown.
